# Real-Time Structural Biology of DNA and DNA-Protein Complexes on an Optical Microscope

**DOI:** 10.1101/2023.11.21.567962

**Authors:** Alan M. Szalai, Giovanni Ferrari, Lars Richter, Jakob Hartmann, Merve-Zeynep Kesici, Bosong Ji, Kush Coshic, Annika Jaeger, Aleksei Aksimentiev, Ingrid Tessmer, Izabela Kamińska, Andrés M. Vera, Philip Tinnefeld

## Abstract

The intricate interplay between DNA and proteins is key for biological functions such as DNA replication, transcription, and repair. To better understand these interactions, it is crucial to develop tools to study DNA-protein complexes with high spatial and temporal resolution. Here, we use the vertical orientation that DNA adopts on graphene and investigate its interactions with proteins via energy transfer from a probe dye to graphene, achieving spatial resolution down to the Ångström scale. We measured the bending angle of DNA induced by adenine tracts, bulges, abasic sites and the binding of Escherichia *coli* endonuclease IV with unprecedented precision and millisecond time resolution. Additionally, we observed the translocation of the O^6^-alkylguanine DNA alkyltransferase along double-stranded DNA, reaching single-base pair resolution and detecting an affinity for adenine tracts. Overall, we foresee that this method will become a widespread tool for the dynamical study of nucleic acid and nucleic acid-protein interactions.

## Main

DNA serves as the repository of information for protein synthesis and the development of cellular structures. The successful storage and use of this information relies on the interactions between DNA and a multitude of proteins. The conformation of DNA, including its bending, plays a critical role in these interactions.^1–5^

Among the techniques used to study DNA-protein complexes, Förster Resonance Energy Transfer (FRET) stands out for being non-invasive and providing insights into dynamics at the nanometer scale. Especially, single-molecule FRET (smFRET) has been pushing dynamic structural biology.^6–10^ However, its performance is hindered by the influence of photophysics from both donor and acceptor molecules, and its dynamic range is limited to distances of 3-10 nm.^11,12^ Furthermore, when FRET is employed to assess DNA bending, additional rotations and translations of the double-helix can affect the donor-acceptor distance.^13^

In this work, we present a new tool to study DNA-protein interactions that overcomes these limitations: Graphene Energy Transfer with vertical Nucleic Acids (GETvNA). On the one hand, this method harnesses the d^-4^ distance dependence of the energy transfer from fluorescent dyes to graphene, which has proved to offer sub-nanometer spatial resolution, while surpassing the dynamic range of FRET by fivefold.^14–18^ On the other hand, GETvNA exploits the vertical orientation adopted by a double-stranded DNA (dsDNA) segment immobilized on graphene *via* a single stranded DNA (ssDNA) overhang. We took advantage of this combination to determine at the single-molecule level bending angles of dsDNA containing bulges, which, in a biological cell, form during DNA replication, and are potentially mutagenic if unresolved by the cellular machinery.^19,20^ We also measured small bending angles in DNA bearing adenine tracts (A-tracts), which consist of sequences of at least four consecutive adenines and are known to bend DNA, playing a key role in DNA packaging and DNA-protein affinity.^21–24^ Furthermore, we unveiled the bending angle distribution induced by an abasic site and the further conformational changes induced by the binding of *Escherichia coli* endonuclease IV (Endo IV), a DNA repair enzyme specific for apurinic and apyrimidinic (AP) sites in DNA.^25^ Finally, we demonstrate the potential of GETvNA for real-time tracking of protein diffusion along DNA by measuring the location of the clinically relevant O^6^-alkylguanine DNA alkyltransferase (AGT) with Ångström spatial resolution.

### Orientation on Graphene

GETvNA is based on immobilizing a hybrid DNA construct consisting of dsDNA with an overhanging ssDNA on graphene (Fig. 1a). The complex is adsorbed on graphene due to π-stacking between the bases of the ssDNA overhang and the hexagonal carbon lattice. In the dsDNA part, only the initial base pair is available for this interaction, while the rest is compromised in base stacking. To study the orientation of the dsDNA segment in this type of construct, we prepared samples with dsDNA of different lengths. Each DNA construct was functionalized with a fluorescent dye at the DNA end that is not attached to the graphene surface, as depicted in Fig. 1a (see sequences in Table S1 and sample preparation protocols in Methods and Fig. S1). We performed single-molecule fluorescence lifetime measurements and obtained the distance *z* between the dye and graphene (hereafter also referred to as the height of the dye), using Eqn. 1:

**Figure 1.**
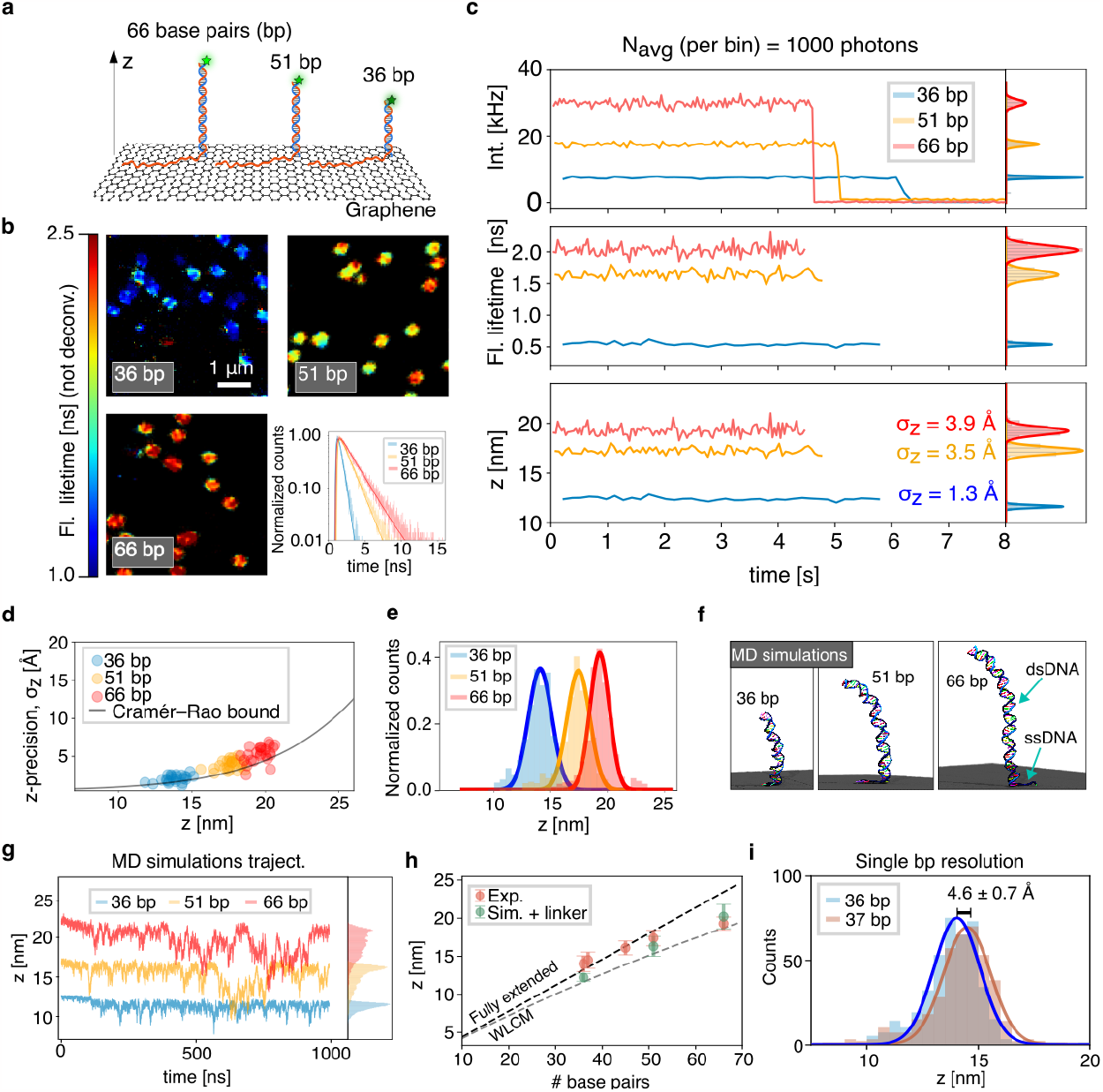
GETvNA principles: dsDNA orientation on graphene determined with Ångström resolution. **a)** Schematic representation of the immobilization of a ssDNA-dsDNA hybrid construct on graphene. dsDNA segments of 36, 51, and 66 base pairs (bp) are shown. Green stars represent the fluorescent label (ATTO 542). **b)** Exemplary FLIM images and single-molecule fluorescence lifetime decays for the three DNA lengths shown in a). **c)** Time traces of representative single molecules. The time bins were adapted to include on average 1000 photons. The axial localization precision reported in the bottom panel was obtained from the standard deviation of the Gaussian fits (right subpanel). **d)** Axial localization precision calculated from 25 single molecule time traces per sample. The solid line shows the Cramér–Rao bound for N = 1000 and SBR_z→∞_ = 75. **e)** Histograms of fluorophores’ height for the three samples described in a). 411, 526, and 475 molecules were analyzed for the 36, 51, and 66 bp samples, respectively. **f)** MD simulation snapshots of the 36 bp (time = 230 ns), 51 bp (time = 990 ns), and 66 bp (time = 1100 ns) systems. **g)** MD simulations trajectories obtained for the three samples including the respective histograms. **h)** Comparison of the experimentally obtained heights with the simulation results, where the contribution of the six-carbon-atom linker was considered. Experimental data as summarized in e), including additional systems of 37 bp (402 molecules) and 45 bp (615 molecules). The 36, 51, and 66 bp systems were simulated once. Experimental heights and error bars were obtained from the mean and the standard deviation of Gaussian fits, while for simulations they were calculated as the arithmetic mean and standard deviation of the height in the trajectory frames.

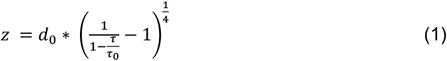

Here, *τ* and *τ*_0_ represent the fluorescence lifetimes in the presence and absence of graphene, respectively, and *d*_0_ is the distance of 50% energy transfer efficiency to graphene (for the fluorophore ATTO 542 we used *d*_0_ = 17.7 nm; see Table S2).^14^

In Fig. 1b, we present FLIM (Fluorescence Lifetime Imaging Microscopy) images obtained for three samples with dsDNA segments of different lengths, where sparse single molecules with homogeneous fluorescence lifetimes are observed (see Fig. S2 for extended fields of view). Additionally, fluorescence lifetime decays of exemplary molecules are depicted. Next, we measured time traces of individual molecules for each of the three samples. In Fig. 1c, we show exemplary time traces of fluorescence lifetime, intensity, and distance to graphene (additional examples are shown in Fig. S3-5). The precision σ_z_ to determine the distance between the dye and graphene was calculated for each time trace, adjusting the time bins for an average of 1000 photons. As highlighted in the bottom panel of Fig. 1c, the precision lies below 4 Å for all cases. In Fig. 1d, we show how σ_z_ varies with *z* for 25 molecules per sample, along with the theoretical limit given by the Cramér–Rao bound (Methods and Fig. S6).

For the three systems, we next calculated the fluorescence lifetime of single molecules using all the photons detected before photobleaching and converted these values into *z* using Eqn.1. In Fig. 1e, the histograms of the obtained distances are depicted. The widths of these distributions lie between 8.5 Å and 10.5 Å, which represent a broader range compared to the precision achieved for individual molecules. This discrepancy is explained by the fact that in a single trace the photon count is solely shot-noise limited, while the histogram of an ensemble of single molecules accounts for local heterogeneity that could e.g. be induced by graphene quality (Supplementary Text and Fig. S7).

As a guide, the expected heights according to a fully extended DNA chain (black dotted line) and the worm-like chain model (WLCM, gray dashed line) are shown. **i)** Height histograms for the 36 bp and 37 bp systems. The mean values of the Gaussian fits were used to calculate the difference in height, while their standard errors were used to estimate the respective error.

The experimental heights retrieved from the Gaussian fits (Fig. 1e) are 14.0, 17.4, and 19.3 nm for the 36, 51, and 66 bp systems, respectively. To complement our experimental data, we performed MD simulations, which strongly suggest that the lowest base pair of the dsDNA segment dictates a vertical orientation on graphene with thermal fluctuations representative for the mechanical properties of dsDNA (see snapshot in Fig. 1f, Fig. S8 and Movies S1-3). In the simulations, the average distances between the last base pair and graphene are 11.2, 15.3, and 19.3 nm (Fig. 1g), following a similar trend as shown experimentally. Interestingly, fast dynamics can be observed in the simulation trajectories, which are averaged out within the integration time of the experimental time traces. However, the fast dynamics become visible on the sub-microsecond timescale when performing a fluorescence autocorrelation analysis (Fig. S9).

The difference in height between experiments and simulations is partially explained by the length of the six-carbon-atom linker that connects the negatively charged dye ATTO542 with the DNA molecule.^2,26^ To determine the position of the dye attached to the end of dsDNA experimentally, we performed measurements on a construct containing 45 bp labeled at one of its end bases, and on another one with 66 bp, internally labeled at base #45. As shown in Fig S10, the difference in height of the dye between these two scenarios is compatible with having an extended linker pointing upwards in the end-labeled case (Fig. S10 and Methods). The heights obtained accounting for this linker are plotted in Fig. 1h, alongside our experimental results. Apart from the aforementioned systems, constructs containing 37 and 45 bp dsDNA segments were also studied (Fig. 1h). Importantly, our data consistently describe the dsDNA segment standing perpendicular to graphene. While the observed heights are close to the contour lengths (black dotted line) for dsDNA of up to 45 bp, this value starts to deviate for longer segments, following a similar trend as described by the worm-like chain model using a persistence length of 50 nm (gray dotted line). The clear difference of the distributions for the 36 bp and 37 bp systems demonstrates single-nucleotide resolution of the technique (Fig. 1i).^27^

### Determining Bending Angles

We first tested our method by measuring bending angles of DNA containing bulges (short unpaired regions). In Fig. 2a, we show a sketch of our experimental design. The bending angle *θ* is calculated from the height of the dye through Eqn. 2:

**Figure 2.**
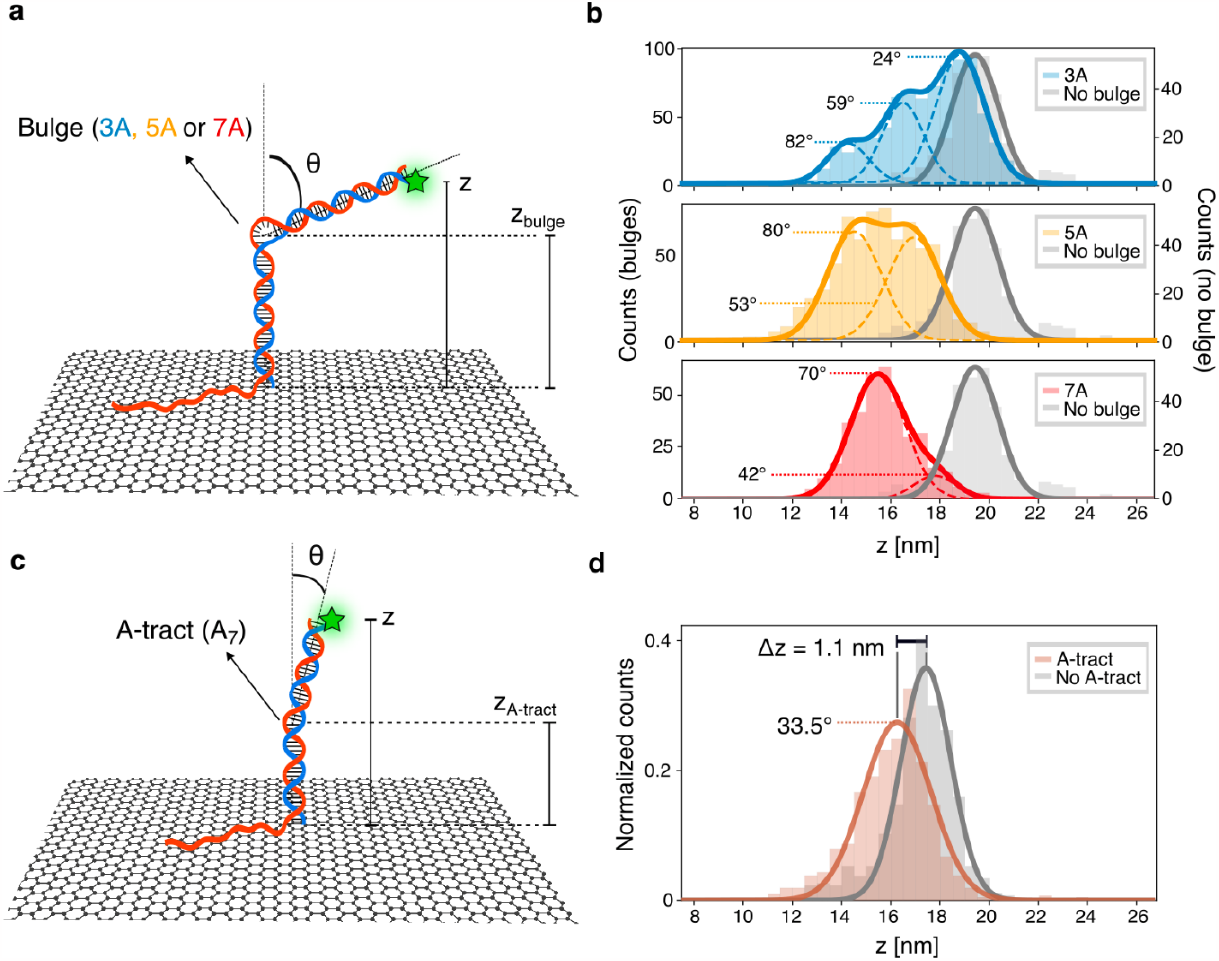
DNA bending originating from bulges and A-tracts. **a)** Sketch of a ssDNA-dsDNA construct immobilized on graphene, containing a bulge in the dsDNA segment. The bending angle (*θ*), fluorophore height (z) and bulge height (*z*_*bulge*_) are highlighted. The green star represents the fluorescent label (ATTO 542). **b)** Height histograms obtained for the three studied systems (858, 705 and 387 molecules for 3A, 5A, and 7A, respectively), alongside the histograms of the analogue dsDNA free of bulges (gray histograms, 475 molecules). Multi-peak Gaussian fits are shown with solid-colored lines, while the individual components of the fits are plotted as dotted lines (see Methods for discussion of the boundaries used in the fits). The bending angles calculated for each subpopulation are presented. **c)** Analogue sketch as in a), with the dsDNA segment containing an A-tract composed of seven adenines. **d)** Height histograms of the sample with (836 molecules) and without (526 molecules) A-tract.

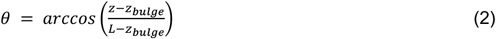

Here, *z* and *z*_*bulge*_ represent the dye-graphene and bulge-graphene distances, respectively, and *L* denotes the height attained by the analogous dsDNA sequence that is free of bulges. The values used for a construct containing a dsDNA with 66 bp and a bulge centered at bp #37 (counting from graphene) were *L* = 19.3 nm and *z*_*bulge*_ = 13.4 mn (Methods). It is noteworthy that *θ* does not depend on *d*_0_ as all three experimental heights from Eqn. 2 are linearly dependent on this parameter.

We studied bulges consisting of 3, 5 and 7 adenines (hereafter called 3A, 5A, and 7A). In previous FRET studies, broad ranges were reported due to multiple possible conformations originating from rotations and translations of the dsDNA segments.^28,29^ Woźniak and co-workers performed twelve measurements labeling different bases close the bulge to get one bending angle.^13^ In contrast, due to the planar geometry of graphene and the distance range of the energy transfer process, GETvNA is mainly sensitive to DNA bending (Supplementary Text and Fig. S11). Therefore, GETvNA directly provides angles according to the simple model represented by Eqn. 2 and Fig. 2a.

In Fig. 2b, the height histograms obtained for the three samples are depicted. By computing the average distances to graphene for 3A and 5A, we retrieved average bending angles of 47° and 69°, respectively. These values are compatible with those reported by Woźniak et al. from smFRET measurements (56° and 73°) and by Dornberger et al. from NMR studies (73° for A5).^13,30^ However, a closer look reveals that 3A exhibits three subpopulations, centered at *z* = 18.7 nm (*θ* = 24°), 16.5 nm (59°), and 14.6 nm (82°). Our findings are in good agreement with an X-ray interferometric study complemented with MD simulations, where the authors proposed that a 3A system is composed of an ensemble of structures with bending angles ranging from 24° to 85°.^31^ For 5A and 7A, the presence of subpopulations is less evident. However, since the distributions are substantially broader than for the straight dsDNA (gray), it can be concluded that multiple conformations with different bending angles coexist at room temperature. A two-peak Gaussian fit unveils center positions at *z* = 14.5 nm (*θ* = 80°) and 16.9 nm (53°) for 5A, as well as 15.5 nm (70°) and 17.8 nm (42°) for 7A. We performed a fluorescence autocorrelation analysis for ten exemplary time traces for the three samples carrying bulges and the one without a bulge. Interestingly, no conversion between states or any other dynamic process were detected on a timescale ranging from 10 μs to 1 s (Fig. S12). In Fig. S13, exemplary time traces of each bulged system are shown alongside their respective fluorescence decay curves. Overall, the lack of dynamics above 10 μs and the monoexponential behavior of the fluorescence decay evidence that the states with different degree of bending were stable at room temperature, only showing subtle dynamics at the (sub-) microsecond timescale.

Next, we used GETvNA to assess the bending angle of an A-tract. For this purpose, we used a sequence containing 51 bp that included an A-tract of seven consecutive adenines (A_7_) centered at 27 bp from graphene, as depicted in Fig. 2c. To calculate the bending angle, we used *L* = 17.4 nm and replaced *z*_*bulge*_ by *z*_*A–tract*_= 10.3 nm in Eqn. 2 (Methods and Fig. S14). Exemplary time traces are depicted in Fig. S15. As shown in Fig. 2d, we obtained a bending angle of 33.5° ± 0.8°, which is slightly larger than the 19°-22° reported for a A_6_ tract,^32,33^ demonstrating that GETvNA allows us to determine small angular deviations.

### Enzyme-Induced DNA Bending

To study the bending of DNA induced by the binding of a cleavage-inactive mutant of *E coli’s* AP Endonuclease IV (Endo IV), a key enzyme of the base excision repair system (BER),^34,35^ we prepared a sample containing an AP site and a phosphorothioate (PTO) modification in the dsDNA segment (see Fig. 3a and Fig. S16). The latter was included as a second safeguard to avoid DNA cleavage by the enzyme. As a control, we first studied the system with a dsDNA segment either carrying or lacking the PTO modification by smFRET. In both cases, after adding Endo IV, we observed the same two FRET states, the one with higher FRET efficiency representing the case where the enzyme bends the AP-bearing DNA as reported by structural studies. ^34,35^ While the PTO modification did not alter the recognition by the enzyme, it increased the frequency of transition between the two states (Figure S17). Therefore, we decided to continue our studies with the DNA containing the PTO modification.

**Figure 3.**
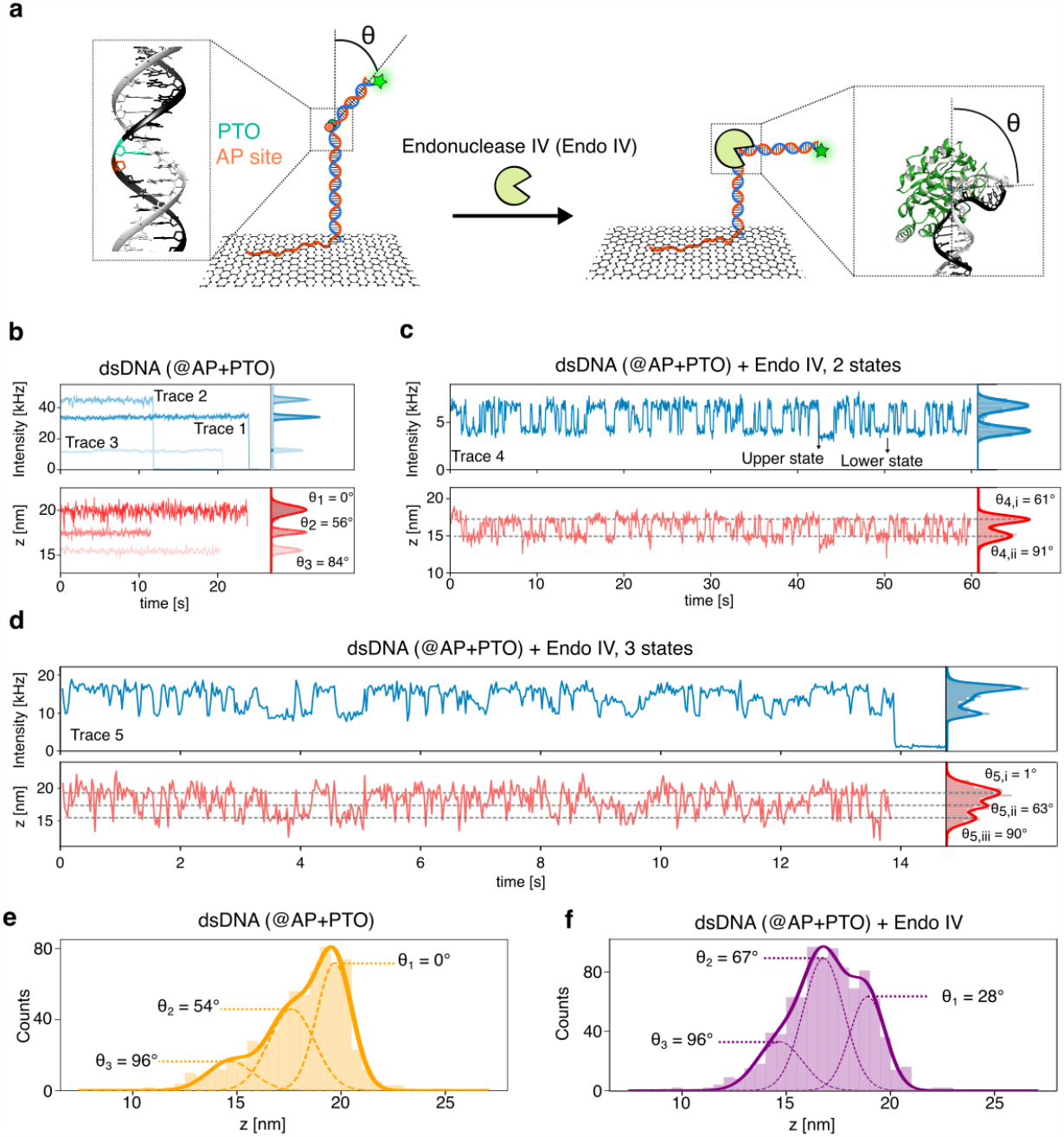
DNA bending induced by Endo IV. **a)** Sketch of the ssDNA-dsDNA containing an AP site and a PTO modification immobilized on graphene, before (left) and after (right) the binding of Endo IV. A B-DNA structure with an AP site labeled in orange and a PTO in green is shown on the left, and the Protein Data Bank structure of Endo IV bound to DNA (2NQJ) is depicted on the right.^34^ The green star from the sketches represents the fluorescent label (ATTO 542). **b)** Exemplary time traces obtained in the absence of Endo IV. On the right, Gaussian fits are presented for each trace alongside the bending angles obtained from the mean heights. **c)** Time trace acquired after adding Endo IV, where two states are resolved. An upper and lower state are defined in the intensity trace. Two-peak Gaussian fits are shown in the right panels, as well as the corresponding bending angles. The gray dotted lines in the height trace indicate the mean heights obtained from the Gaussian fits. **d)** Similar time traces as in e) showcasing three states. **e)** Height histogram for traces acquired without Endo IV in solution (667 molecules). The distribution is well-described by the sum of three Gaussian functions, which are shown as dotted lines. The bending angles associated with each subpopulation are shown. **f)** Height histogram including all states detected in traces showing state changes in the presence of Endo IV. The three-peak Gaussian fit is shown as a solid line, while its individual components are displayed as dotted lines.

We performed GETvNA measurements in the presence and in the absence of Endo IV. We retrieved bending angles using *z*_*AP*_ = 15.1 nm instead of *z*_*bulge*_ in Eqn. 2 (see Methods). In Fig. 3b, three exemplary time traces without enzyme are shown. Here, we measured stable heights of 15.5 nm, 17.5 nm, and 20.0 nm, corresponding to bending angles of 84°, 56° and 0°, respectively. It is important to highlight that 0° refers to the absence of bending in comparison to the unmodified, straight dsDNA. After analyzing 667 molecules, we obtained three populations, one of them reproducing the height of dsDNA without the AP site (0°), another one being centered at *θ* = 54°, and a minor population with *θ* = 96°, as depicted in Fig. 3e (additional exemplary time traces are shown in Figure S18). These observations suggest that an AP site can introduce bent conformations due to the interruption of the π-stacking between consecutive bases at the abasic site. Yet, the structure of a fraction of molecules remains unaffected and keeps the conformation of the dsDNA without AP site and PTO modification. To compare this result with previous reports, we analyzed the conformation of AP-bearing DNA molecules previously obtained by NMR and found bending angles in the 43-53° range (Fig. S19), which agree with the 45° obtained from our average height of 18.1 nm.^36^

After adding Endo IV to the solution, we observed a similar behavior as in our smFRET studies. As exemplified in Figs. 3c and 3d, our GETvNA measurements showed dynamic switching between states with different degrees of bending due to the action of Endo IV. Fig. 3c shows a two-state single-molecule trace (*θ* = 61° and 91°), while the example from Fig. 3d corresponds to a three-state system (*θ* = 1°, 63°, and 90°). More examples of traces showing dynamic behavior are shown in Fig. S20. Notably, GETvNA resolves transitions occurring in the tens of millisecond timescale between states that differ in their bending angles by less than 30°.

Finally, in Fig. 3f we show the distribution of bending angles obtained from 394 time traces showing state changes. These dynamic cases represent 40% of a total of 993 acquired traces (see Fig. S21 and Fig S22). Interestingly, the distribution can be described by the sum of three Gaussian functions, representing stable conformations with *θ* = 28°, 67° and 96° and *σ*_*θ*_ = 24°, 15°, and 15°, respectively. On the one hand, these values were the same as those obtained from the static traces in the presence of Endo IV (Fig. S21), suggesting that the actual stable conformations were the same independent of the switching behavior. On the other hand, they were almost identical to the angles retrieved from traces where the dsDNA lacked a PTO modification (Fig. S23), reproducing our smFRET observations. Notably, the mean values of these distributions were also similar to those obtained in the absence of Endo IV, but with the state with intermediate bending being visited with higher frequency when the enzyme was present. Additionally, the peaks corresponding to the intermediate and least bent states are slightly shifted towards higher angles, which can be related to a charge screening effect given the positive charge of Endo IV, which may allow for shorter distances between the DNA on both sides of the bound protein. Strikingly, the bending angle of the main population was 67°, which lies close to the 74° that we extracted from the crystallographic structures available (Fig. S24 and Methods). ^34,35^ The interplay between three states with different degrees of bending was previously observed for other proteins,^37,38^ but we report this behavior for Endo IV for the first time. Additionally, we provide the actual bending angle of each state at the single-molecule level, something impossible to obtain with smFRET without accounting for the additional rotations and translations that affect the FRET outcome (Supplementary Text and Fig. S11).

### Protein Diffusion

GETvNA can also be applied to directly DNA interacting proteins. To this end, we tracked the diffusion of AGT, a protein that repairs the highly mutagenic and cytotoxic O^6^-alkylguanine lesion by performing the tasks of lesion search, identification, and removal.^39,40^ The employed repair-inactive mutant of the human protein was labeled with ATTO 647N, as sketched in Fig. 4a (purple star). In Fig. 4b, we present an exemplary time trace with 50 ms time bins, where bidirectional movement is observed. In the zoomed-in area displayed in Fig. 4c, a fast and a slow mode can be discerned as recently reported.^41^ However, previous studies were not able to resolve the translocation of AGT in the slow mode. Here, the Ångström precision achieved by GETvNA enables us to observe steps, as depicted in Fig. 4c, where a 7.0 Å displacement is highlighted. A similar diffusive behavior was observed for AGT clusters, as exhibited in Fig. S25 (see Methods for more details).

**Figure 4.**
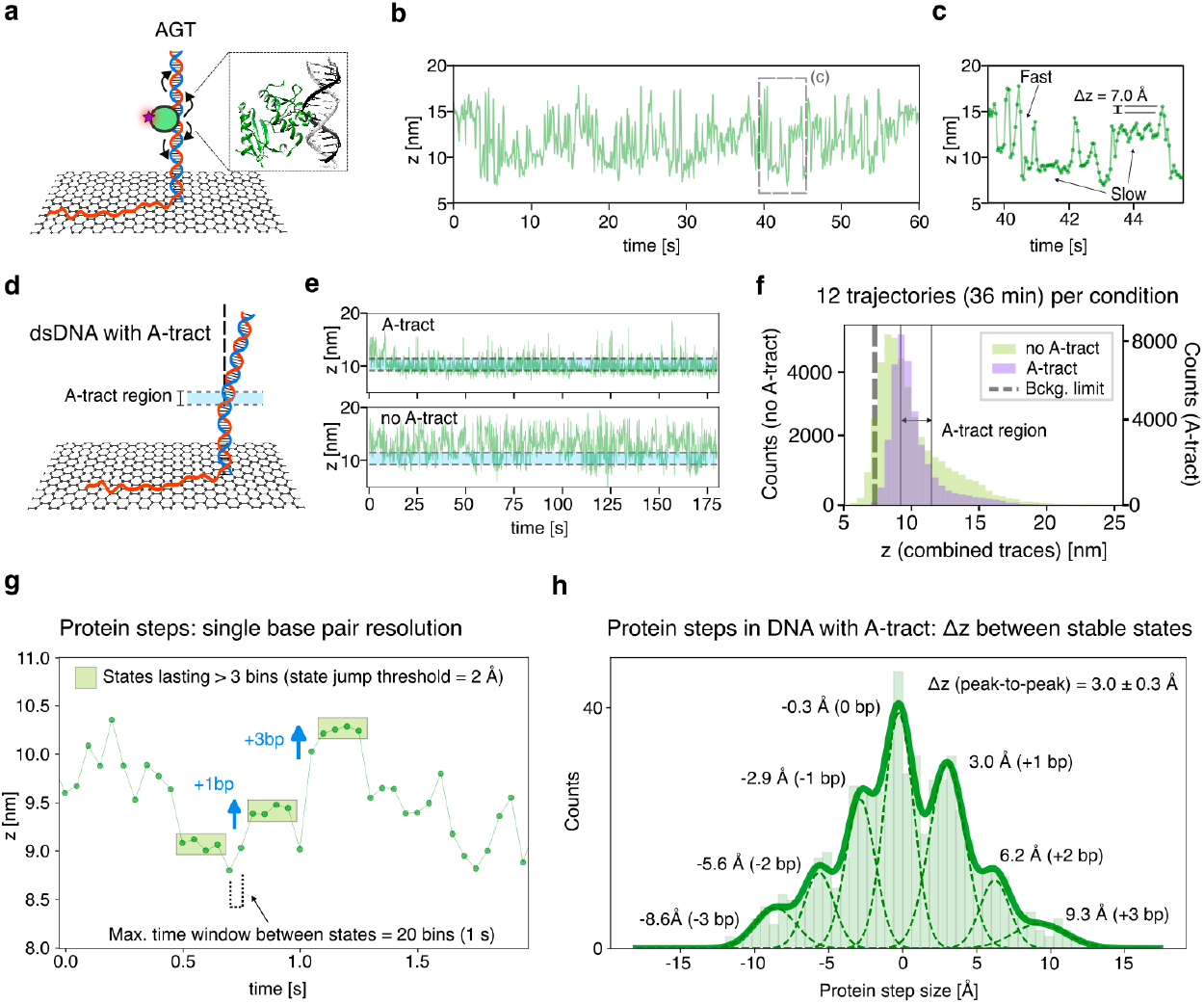
AGT diffusion on dsDNA. **a)** Sketch of a ssDNA-dsDNA construct immobilized on graphene with an AGT protein bound to it. Curved arrows represent the possible protein diffusion and the purple star the fluorescent label (ATTO 647N). The Protein Data Bank structure of AGT bound to DNA (1YFH) is depicted on the right.^39^ **b)** Exemplary time trace of an AGT monomer diffusing on dsDNA. The time binning of this trace (and all traces within this figure) is 50 ms. **c)** Extended view of the time lapse marked in b). Fast and slow modes are highlighted, as well as a protein step of 7.0 Å. **d)** Sketch of a construct with a dsDNA segment containing an A-tract. The axial boundaries of the A-tract are indicated by dotted lines. **e)** Exemplary time traces of AGT monomers diffusing along constructs with and without A-tract. The axial region corresponding to the position of the A-tract is shadowed in light blue and framed by gray dotted lines. **f)** Histograms of heights obtained from twelve time traces (180 seconds per trace) with A-tract and twelve without A-tract. The A-tract region is highlighted (grey vertical lines). The background-limited height is marked with a black dashed line. **g)** Exemplary fragment of a time trace corresponding to an AGT cluster diffusing on DNA with an A-tract, where the filtering step used to quantify protein steps is shown. The identified stable states (at least four consecutive bins where the height difference between adjacent bins is smaller than 2 Å) are highlighted with green rectangles. Blue arrows indicate the magnitude of the step between the marked stable states. **h)** Histogram of protein step size, obtained for all the data acquired on DNA containing an A-tract after performing the filtering step described in g). A seven-peak Gaussian fit is shown as a solid line, while each component of the fit is shown with a dashed line, alongside their respective mean values. The weighted average of peak-to-peak distances is also presented, together with the weighted standard deviation (weights are calculated from the areas under each Gaussian curve).

Next, we evaluated the effect of having a pronounced bent in the dsDNA on the diffusive behavior of AGT monomers, by using dsDNA containing an A_7_ A-tract. Remarkably, we observed a restricted movement around the height where the A-tract was located, as shown in the top trace from Fig. 4e. This behavior contrasts the diffusion observed in the absence of an A-tract, where no preferential residence in this same region was observed (Fig. 4e, bottom). In Fig. 4f, the height histograms obtained from single-molecule time traces with and without A-tract are plotted, showing that the fraction of time spent outside of the *z* = 9.2-11.5 nm region (namely, the region where the A-tract is located) was lower for traces where the dsDNA contained an A-tract. A similar behavior was observed for AGT clusters, as depicted in Fig. S26. In Fig. S27, we show that the fraction of time spent between *z* = 9.2-11.5 nm was 49% (10% standard deviation, SD) for monomers and 58% (7% SD) for clusters when the A-tract was present, and 28% (15% SD) and 24% (17% SD) when there was no A-tract. It is worth noting that heights below 7 nm were not accessible, because the fluorescence quenching was above 98% (see Fig. S28). In Fig. S29, additional exemplary traces of monomers and clusters in the presence and absence of an A-tract are presented.

As discussed above, our GETvNA measurements allowed detecting sub-nanometer changes in the position of AGT in its slow mode. As the movement observed on dsDNA containing an A-tract was spatially restricted, we evaluated the slow diffusion in those cases, to study if such movement corresponded to stepwise translocation. For this purpose, we kept only bins describing long-lived states, the position of which could be determined with high accuracy (Supplementary Text and Fig. S30). In Fig. 4g, an exemplary fragment of a time trace is shown, where the detected stable states are highlighted. The protein steps were quantified as the difference in height between consecutive stable states. The histogram of protein steps shown in Fig. 4h exhibits peaks that are separated by 3.0 ± 0.3 Å, in agreement with the expected values for -3, -2, 1, 0, 1, 2, and 3 base pairs steps. The inter-base pair distance of 3.0 ± 0.3 Å is in excellent agreement with the 3.1 Å obtained from the *z*-projection of a 3.4 Å vector tilted by ∼20°, which is close to average tilting observed for bp #31-36 segment (see Fig. S8d and Supplementary Text). The zero-base-pair peak appears due to the temporal window introduced in the filtering step, which allows detecting these transitions whenever the protein returns to the same base pair after visiting a neighboring one for a short period. In Figure S31, we show the protein step size histogram for the data acquired on DNA lacking an A-tract, where we still observe a distribution describing jumps from -3 to 3 base pairs with 3.0 ± 0.7 Å inter-base pair distance. In summary, our results show that, in its slow mode, AGT moves stepwise on DNA with minimal steps of single base pairs, demonstrating that GETvNA achieves single-base pair resolution to track protein motion on DNA.

## Discussion and Outlook

We presented GETvNA, a robust platform to study DNA-protein interactions and conformational changes of DNA with Ångström resolution on a confocal microscope. We studied different biologically relevant DNA bending scenarios, such as DNA containing A-tracts or bulges, as well as bending induced by an enzyme. While our results agree with previous studies, we were additionally able to resolve intermediate states and observe dynamic behaviors in ambient conditions at the single-molecule level. As shown here, this can be particularly useful when studying the DNA repair machinery, where GETvNA unveils the dynamic dimension, thus complementing the static structural information available by high resolution structural methods. We showed that GETvNA can outperform smFRET in terms of accuracy and dynamic range, while maintaining a straightforward workflow. In addition, DNA conformational heterogeneity is readily accessible with GETvNA. By requiring labeling with only a donor dye, the GETvNA outcome is not affected by acceptor photophysics and hence has a simpler interpretation compared to FRET as well as interference with biomolecular activity is less likely. Finally, dye-labeled molecules unspecifically binding to graphene are fully quenched reducing background contributions.

To date, fluorescence-based methods reaching Ångström resolution were only demonstrated in techniques requiring the accumulation of multiple localizations within minutes, therefore lacking temporal resolution.^42,43^ Regarding tracking applications, MINFLUX has recently been used to track the one-dimensional diffusion of motor proteins on microtubules, being able to resolve 4 nm sub-steps.^44,45^ Here, we demonstrated that GETvNA allows tracking the movement of proteins on DNA by observing the diffusion of AGT. We achieved a spatial resolution of ∼3 Å that enabled the detection of single-base pair displacements, something that typically needs measuring forces with complex optical tweezers setups,^46–49^ and here it was reached using a standard confocal microscope.

The inherent simplicity of GETvNA, in combination with its exceptional spatial resolution, sets it apart as an exceptional tool for unraveling the intricate conformational details and underlying mechanisms driving the formation of DNA-protein complexes. GETvNA therefore has the potential to become a widespread technique and unlock a world of exciting possibilities in the realm of structural biology. Moreover, the demonstration that dsDNA stands vertically on graphene will pave the way for new orientational studies of DNA on related materials, bringing up thrilling applications in an emerging field that combines single-molecule biophysics with 2D materials.^50^

## Acknowledgements

The authors thank for financial support by the Deutsche Forschungsgemeinschaft (DFG, German Research Foundation) under grant numbers TI 329/14-1 and KA 5449/2-1, the excellence cluster e-conversion under Germany’s Excellence Strategy – EXC 2089/1 – 390776260, and by the Center for NanoScience (CeNS). Furthermore, funded by the Federal Ministry of Education and Research (BMBF) and the Free State of Bavaria under the Excellence Strategy of the Federal Government and the Länder through the ONE MUNICH

Project Munich Multiscale Biofabrication. L.R. acknowledges support by the Studienstiftung des deutschen Volkes. A.M.S. is thankful for the support by the Alexander von Humboldt foundation under reference Ref 3.2 - ARG - 1220722 - GF-P. I.K. acknowledges support by the National Science Center of Poland (Sonata 2019/35/D/ST5/00958). K.C. and A.A. were supported by the US National Science Foundation (DMR-1827346) and the Human Frontier Science Program (RGP0047/2020). The supercomputer time was provided through ACESSS allocation grant MCA05S028 (A.A.) and the Leadership Resource Allocation MCB20012 on Frontera of the Texas Advanced Computing Centre (A.A). I.T. acknowledges financial support by the Deutsche Forschungsgemeinschaft (DFG, German Research Foundation), under grant number TE671/7-1. L.R. acknowledges Stefan Krause who suggested preliminary experiments leading to the discovery of GETvNA. Furthermore, he thanks Patrick Schüler, Tim Schröder, and Jonas Zähringer for fruitful discussions.

## Author contributions

P.T., A.M.S., and L.R. conceived the concept and experiments. A.M.S., G.F., and L.R. designed the experiments and the analysis pipeline, and curated data. A.M.S., G.F., L.R., J.H., M.K., B.J., A.J., A.M.V., and I.K. conducted experiments. A.M.S. and G.F. developed the analysis software. A.M.S., G.F., L.R., J.H., M.K., and B.J. analyzed data. K.C. and A.A. contributed the molecular dynamics simulations. A.M.V. contributed to the design of studies involving Endo IV and their interpretation. I.T. contributed to the design and interpretation of experiments involving AGT and contributed AGT samples. B.J., M.K., and A.M.V. prepared Endo IV samples. I.K. prepared graphene-on-glass samples, optimized their preparation protocol, and interpreted data. P.T. supervised the study. A.M.S. supervised data acquisition, analysis, and visualization. A.M.S., G.F., L.R., and P.T. interpreted data and wrote the manuscript. All authors reviewed and approved the final manuscript.

