## Supplementary material for "Real-Time Structural Biology of DNA and DNA-Protein Complexes on an Optical Microscope": SI_text

Fig. S3. Exemplary time traces of systems containing a 36 bp-long dsDNA segment. .. 29

Fig. S4. Exemplary time traces of systems containing a 51 bp-long dsDNA segment. .. 30

Fig. S5. Exemplary time traces of systems containing a 66 bp-long dsDNA segment. .. 31

### 1 Materials and Methods

If not otherwise stated, laboratory products were purchased from Carl Roth GmbH + Co. KG, Germany and chemicals from Sigma-Aldrich Chemie GmbH, Germany.

#### 1.1 Graphene Preparation

Monolayer graphene on a 60 mm × 40 mm copper substrate with poly(methyl methacrylate) (PMMA) on top was purchased from ACS Materials LLC, USA or Graphenea Inc., Spain. The foil was stored in a vacuum-sealed desiccator. A wet-transfer approach was used to transfer the graphene to glass coverslips (Paul Marienfeld GmbH & Co. KG, Germany)[1]. All coverslips were treated with ultrasound at 37˚C for three times 15 minutes in a bath of Milli-Q water to remove any contaminants from the surface. In the first of the three rounds, 1% v/v of Hellmanex® III (Hellma GmbH & Co. KG, Germany) was added to the bath.

Smaller pieces of roughly 0.25 $cm^2$ were carefully cut from the PMMA/graphene/copper foil. The copper was wet-etched by letting a piece float with the copper film exposed to 0.2 M ammonium persulfate ($(NH_4)_2S_2O_8$) for ~4 h. A coverslip was dipped vertically while slowly moving towards the PMMA/graphene, scooped gently out of the solution, and transferred to Milli-Q water in order to wash out the residues of ammonium persulfate. The step of washing PMMA/graphene was repeated twice with fresh Milli-Q water. Next, the PMMA/graphene was scooped with a glass coverslip, carefully dried using a nitrogen stream and stored overnight. Extra PMMA ($M_w$ = 120,000 g/mol) was dissolved in chlorobenzene (50 mg/mL) and was drop-casted on top of the first PMMA/graphene layer. This allowed the dried PMMA to re-dissolve, thus relaxing the underlying graphene monolayer and forming an improved contact with the substrate[2]. The added PMMA dried for at least 30 minutes. The PMMA/graphene was directly cleaned as outlined below or stored in Milli-Q water until cleaning and subsequent usage.

For cleaning, the PMMA/graphene on glass was first dipped in two distinct acetone baths for 7 min each. Subsequently, the washing was repeated in toluene for 7 min. After each washing step, the samples were dried with a nitrogen stream. Finally, the graphene-on-glass was placed on active coal that was heated on a heating plate to 230 ˚C. Here, the graphene faces the active coal. The final heating was done for about 10 h prior to usage. The graphene-on-glass is ready to use after the heating.

The whole procedure for transferring and cleaning of graphene to glass coverslips is depicted in Fig. S1.

1.2 Chemicals/ Buffers

| **24% denaturing acrylamide stock** | |
|---|---|
| Tris | 90 mM |
| Boric acid | 90 mM |
| Acrylamide | 24% v/v |
| Urea | 7 M |

| **10× TBE buffer** | |
|---|---|
| Tris | 900 mM |
| Boric acid | 900 mM |
| EDTA | 11 mM |

| **Elution buffer** | |
|---|---|
| Ammonium acetate | 500 mM |
| Magnesium acetate | 10 mM |

| **1× TAE buffer** | |
|---|---|
| Tris | 40 mM |
| Acetic acid | 20 mM |
| EDTA | 1 mM |
| Sodium chloride | 500 mM |

| **Tx-Tq-PCA buffer** | |
|---|---|
| Tris | 40 mM |
| Acetic acid | 20 mM |
| EDTA | 1 mM |
| Sodium chloride | 500 mM |
| Trolox/Troloxquinone (Tx-Tq) | 2 mM |
| Methanol (optional, for dissolving Tx-Tq) | 0.5 mM |

| | |
|---|---|
| 3,4-Dihydroxybenzoic acid (PCA) | 12 mM |

| 50× PCD buffer | |
|---|---|
| PCD (protocatechuate 3,4-dioxygenase from *pseudomonas sp.*) | 2.8 mM |
| Glycerol | 50% v/v |
| KCl | 50 mM |
| Tris HCl | 100 mM |
| EDTA-$Na_2 \cdot 2H_2O$ | 1 mM |

| NEB3-Tx-Tq-PCA buffer (pH 7.9) | |
|---|---|
| Sodium chloride | 100 mM |
| Magnesium chloride | 10 mM |
| Tris | 50 mM |
| Trolox/Troloxquinone (Tx-Tq) | 2 mM |
| 3,4-Dihydroxybenzoic acid (PCA) | 12 mM |

| AGT labeling buffer | |
|---|---|
| 4-(2-hydroxyethyl)-1-piperazineethanesulfonic acid (HEPES; pH 7.5) | 25 mM |
| Sodium acetate | 25 mM |
| Magnesium acetate | 10 mM |

| AGT experiment buffer | |
|---|---|
| Tris-HCl (pH 7.9) | 10 mM |
| Sodium chloride | 50 mM |
| Dithiothreitol (DTT) | 1 mM |

#### 1.3 DNA Preparation

DNA oligonucleotides were purchased from biomers.net GmbH, Germany (Biomers); Eurofins Genomics Europe Shared Services GmbH, Germany (Eurofins); and Integrated DNA Technologies, Inc., USA (IDT). See Table S1 for a full list of the used oligonucleotides and their respective supplier.

##### 1.3.1 Purification

Single-stranded DNA (ssDNA) strands of more than 50 nt length were purified on 8 M urea polyacrylamide gel (PAGE) for 1 h (100 V, 15-8 mA) using the 1× TBE buffer. After electrophoresis, the gels were illuminated with a UV lamp at 254 nm (Analytik Jena GmbH + Co. KG, Germany) or a blue LED transilluminator (IO Rodeo, US) at 470 nm and the bands were excised. The gel pieces were cut and squeezed, incubated in 400 $\mu$L of the elution buffer, and frozen at –20 °C. The solution was shaken for 3 h and the gel debris was separated with a Spin-X filter (Favorgen Biotech Corp., Austria) for 2 min at 10,000 g, rt. 100 µL of the elution buffer was added to the gel debris and centrifuged again at 10,000 g for 2 min. The DNA strands were desalted by dialysis. The sample was concentrated with an 3K Amicon Ultra Filter (Merck Millipore, Merck KGaA, Germany) to a concentration of 1 µM or higher. DNA was quantified by measuring UV absorbance at 260 nm or the absorbance of the specific dye with a photospectrometer (Thermo Scientific NanoDrop 2000, Thermo Fisher Scientific Inc., USA). Samples were loaded onto a polyacrylamide gel in 1× TBE buffer (60 min at 100 V) and stained with Sybr™ Gold (invitrogen, Thermo Fisher Scientific Inc., USA) for final visualization of the purified DNA strands.

During the whole purification, ssDNA strands with an attached fluorescent dye were handled separately from those without.

##### 1.3.2 Annealing for GETvNA

For graphene energy transfer with vertical nucleic acid (GETvNA) experiments, ssDNA strands were annealed to a respective reverse complementary strand by mixing both in 1× TAE buffer solution. The mixing was performed having a 10× excess of the longer ssDNA strand that served as a toehold on graphene. After premixing, the solution was shaken at 300 rpm and at 37 °C on a thermomixer (Eppendorf ThermoMixer® C, Germany) for 2 h to induce the formation of a double-stranded DNA (dsDNA).

##### 1.3.3 Annealing for Single-Molecule FRET

For single-molecule (smFRET) experiments, four strands had to be annealed to form a complex. Namely, the red dye-labeled strand, the green dye-labeled strand, the reverse complementary strand with biotin, and the AP strand with or without PTO modification were annealed. The combined strands were mixed at the same concentration in NEB3 buffer without photostabilizing agents. The solution was heated to 90 °C before cooling to 24 °C, while having a temperature drop of 1 K/min, on a thermocycler (eppendorf Mastercycler®).

#### 1.4 Enzyme Preparation

##### 1.4.1 Endonuclease IV

The endonuclease IV gene from *Escherichia coli* was PCR amplified from chromosomal DNA and cloned between the NcoI XhoI restriction sites of the expression plasmid pET24-d (Novagen, Merck Millipore, Germany). The E261Q endonuclease IV activity mutant[3] was created by overlapping PCR and cloned in the pET24-d vector using the same restriction sites. All enzymes for cloning were purchased from New England Biolabs (USA) and the *E.coli* XL1blue strain was used for cloning.

The E261Q endonuclease IV mutant was produced in the *E. coli* expression strain BL21 Star (DE3). The cells were grown in LB supplemented with 30 µg/ml of kanamycin at 37 °C. When the cell culture reached an $OD_{600}$ of 1, the expression was induced with 1 mM IPTG for 4 h at 25 °C. The protein was purified by metal affinity chromatography.[4] After the affinity purification, the sample was further purified by cationic exchange chromatography using HiScreen SP HP columns (Cytiva Europe GmbH, Germany). The cationic exchange chromatography was performed in 20 mM Tris, 100 mM NaCl, 0.2 mM EDTA (pH 7.4) and the protein was eluted from the column using a linear gradient to a 20 mM Tris, 1 M NaCl, 0.2 mM EDTA buffer (pH 7.4). The sample was stored at -20 °C in 50% glycerol 10 mM Tris, 50 mM NaCl, 0.1 mM EDTA (pH 7.4).

##### 1.4.2 AGT

AGT was purified from *E.coli* as previously described[5], and fluorescently labeled with ATTO 647N (ATTO-TEC GmbH, Germany) *via* maleimide linkage. To avoid interference of fluorophore conjugation with the protein's DNA binding interface, we used the (catalytically inactive) $AGT_{C145S}$ variant. Labeling was performed at 10 µM protein concentration and 10-fold excess ATTO 647N for 1 h at ambient temperature in a 100 µl volume, using the AGT labeling buffer. The labeled protein was purified from excess of free dye and transferred to the AGT experiment buffer at 4 °C using a 7 kDa MWCO Zeba spin desalting column (Thermo Fisher Scientific Inc., USA). 1.5:1 dye:protein labeling ratio was confirmed by absorption measurements with a NanoDrop ND1000 spectrophotometer (Peqlab, Germany), using extinction coefficients of 150,000 $M^{-1}cm^{-1}$ for ATTO 647N at 650 nm (correction factor at 280 nm: 0.049), and 26,720 $M^{-1}cm^{-1}$ for AGT at 280 nm (minus correction for ATTO 647N contribution). Functionality of labeled AGT in binding to its target lesion (methylguanine) was confirmed using electrophoretic mobility shift assay (EMSA).

#### 1.5 <u>Setups</u>

##### 1.5.1 Confocal Setup

All experiments involving graphene were performed on a home-built confocal microscope based on an Olympus IX-71 inverted microscope. Here, the sample was excited by pulsed lasers at 532 nm or 636 nm (LDH-P-FA-530B and LDH-D-C-640; both PicoQuant GmbH, Germany) at a frequency of 40 MHz (PDL 828 "Sepia II", PicoQuant GmbH with an oscillator module: SOM 828, PicoQuant GmbH, Germany). The lasers were coupled into a single mode fiber (P3-488PM-FC, Thorlabs GmbH, Germany) to obtain a Gaussian beam profile and to perfectly overlay the two excitation beams. Circular polarized light was obtained by a linear polarizer (LPVISE100-A, Thorlabs GmbH, Germany) and a quarter-wave plate (AQWP05M- 600, Thorlabs GmbH). The light was focused in a diffraction-limited spot using an oil-immersion objective (UPLSAPO100XO, NA 1.40, Olympus Deutschland GmbH, Germany). The position of the sample is adjusted using a piezo stage (P-517.3CD, Physik Instrumente (PI) GmbH & Co. KG, Germany) and a controller (E-727.3CDA, Physik Instrumente (PI) GmbH & Co. KG, Germany). The emission light was separated from the excitation beam by a dichroic beamsplitter (zt532/640rpc, Chroma Technology Corporation, USA) and focused onto a 50 µm diameter pinhole (Thorlabs GmbH, Germany). After the pinhole, signals of different wavelengths were separated by a dichroic beamsplitter (640 LPXR, Chroma Technology Corporation, USA) into a green (Brightline HC582/75, AHF, Germany; RazorEdge LP 532, IDEX Health & Science, LLC, USA) and red (SP 750, AHF, Germany; RazorEdge LP 647, IDEX Health & Science, LLC, USA) detection channel. The emission was focused onto avalanche photodiodes (SPCM-AQRH-14-TR, Excelitas Technologies Corporation, USA) and the signals were registered by a time-correlated single photon counting (TCSPC) unit (HydraHarp400, PicoQuant GmbH, Germany). The setup was controlled by a commercial software package (SymPhoTime64, Picoquant GmbH, Germany).

For measurements on the surface of a graphene sheet, the laser powers in front of the objective were chosen to be 3-4 µW at 532 nm and 4 µW at 640 nm. FLIM maps, as shown in Fig. 1b, were acquired using 50 nm pixel size and a dwell time per pixel of 2 ms. Unquenched fluorescence lifetimes were measured in solution with laser powers in front of the objective set to 3 µW at 532 nm and at 0.3 µW at 640 nm, focusing about 10 µm from the glass surface into the solution.

##### 1.5.2 Widefield Setup

Comparative smFRET experiments were carried out on a commercial Nanoimager S (ONI Ltd., UK) single-molecule widefield fluorescence microscope. The red excitation at 640 nm and the green excitation 532 nm were realized with a laser power of 3.1 mW and 11.8 mW, respectively. The microscope was set to TIRF illumination at an angle of 55° to the optical axis. The emission was imaged in two spectral channels in green and in red to extract ratiometric intensity values. Here, the pixel size was 117 nm.

#### 1.6 Experiments

##### 1.6.1 GETvNA

A chamber (SecureSeal™ hybridization chambers, Grace Bio-Labs, USA) was glued around the sheet of graphene on the glass coverslip. The desired mixture containing the annealed DNA oligonucleotides was incubated for 1 min. The chamber was rinsed with the 1× TAE buffer. For photostabilization (oxygen removal and triplet state suppression), the Tx-Tq-PCA buffer was mixed with the 50× PCD buffer in a ratio of 1:50 and added to the sample solution. The chamber was sealed air-tight with Microtube Tough-Spots® labels or similar adhesive tags.

In the absence of endonuclease IV (Endo IV), the DNA was incubated as described above. The chamber was washed twice with the 1× TAE buffer. The NEB3-Tx-Tq-PCA buffer was mixed with the 50× PCD buffer in a ratio of 1:50 and added in exchange to the buffer solution. The chamber was sealed air-tight with Microtube Tough-Spots® labels or similar adhesive tags.

For measurements involving Endo IV, the enzyme was prepared at 600 nM concentration in the NEB3-Tx-Tq-PCA buffer. The label on top of the chamber was removed and the enzyme solution was added in exchange for the present buffer. The enzyme solution was incubated for 5 min before it was removed and replaced with an enzyme solution with photostabilizer. For this, the NEB3-Tx-Tq-PCA buffer was mixed with the 50× PCD buffer in a ratio of 1:50 and the enzyme was added at 600 nM concentration. The chamber was sealed air-tight with Microtube Tough-Spots® labels or similar adhesive tags. After about 2 h, if the enzyme activity dropped significantly, or the photostabilization showed no remaining effect, the sealing label was removed and the old, enzyme-containing solution was replaced with new one prior to repeated sealing.

For experiments with the AGT enzyme, the sample with the prepared DNA (66 base pair (bp) dsDNA segment) was incubated with 1 nM BSA for 10 min. The dsDNA probe was functionalized with Cy3B, which was used as a reference to confirm by colocalization that the proteins were bound to dsDNA (AGT was labeled with ATTO 647N). Labeled AGT was added at a concentration of 300 nM leading to the formation of AGT monomers[5]. The labeled AGT was incubated for 2 min before washing the sample 12 times with the AGT experiment buffer, which was kept in the sample for measurements. After washing out unbound proteins, we selected colocalized spots and measured time traces on individual proteins, exciting ATTO 647N only. For cluster formation, the sample was then further incubated with 4 µM of unlabeled AGT for 2 min. Prior to the experiments, the solution was exchanged with the AGT experiment buffer.

For the reference measurements of the unquenched fluorescence lifetimes, experiments were performed in solution since on glass the DNA does not stand perpendicular to the surface, which would make the dye stick to the surface and hinder its free rotation. Instead, a chamber was passivated with BSA-Biotin at a concentration of 1 mg/ml for 10 min and then rinsed twice with 1× TAE buffer. For ATTO 542 in Tx-Tq-PCA buffer, the chamber was filled with a 10 nM solution of labeled and previously annealed dsDNA (36mer + biotinylated 36mer, see Table S1). For this, a solution of the Tx-Tq-PCA buffer with the

50× PCD buffer was mixed in a ratio of 50:1. Similarly, a reference measurement was done with the NEB3-Tx-Tq-PCA buffer mixed with the 50× PCD buffer in a ratio of 50:1. Prior to these experiments, the chamber was sealed air-tight with Microtube Tough-Spots® labels or similar adhesive tags.

For the reference measurements with AGT labeled with ATTO 647N and bound to DNA, the chamber was filled with the AGT experiment buffer containing 300 nM of labeled AGT and 3.7 µM of unlabeled AGT, and 10 µM of unlabeled 66 bp long dsDNA (66mer + 136mer; see Table S1) were added. Unlabeled AGT was added in order to enhance the probability of DNA binding because AGT displays a low $K_D$ for DNA binding of only around 1 µM. In addition, the excess of unlabeled protein decreases the probability of DNA-bound clusters containing more than one fluorophore, thus avoiding interactions between dye molecules. Intensity spikes with a distinct fluorescent lifetime were visible in the fluorescent intensity traces. In the absence of DNA in the chamber, a lower signal was detected likely caused by the freely diffusing proteins as well as dye-labeled proteins sticking to the glass surface in spite of passivation. Without DNA, a fluorescent lifetime of 3.58 ns was measured. Fluorescent signals in the presence of DNA were hence interpreted as slowly diffusing AGT clusters bound to DNA. The reference lifetime for the system was then obtained by fitting only the photons within the intensity spikes for protein-DNA samples. We further filtered out spikes with intensities suggesting the presence of more than one dye per cluster. Fig. S32 depicts the outlined procedure. The obtained fluorescence lifetime from reference measurements is 3.78 ns (as also listed in Table S2), contrasting the previously mentioned value in the absence of DNA.

##### 1.6.2 Single-Molecule FRET

In order to prepare a slide for smFRET measurements, the glass surface was functionalized with a layer of BSA-Biotin-Neutravidin, such that biotinylated samples can be immobilized on the surface. A glass coverslip was first rinsed with Milli-Q grade water and dried with pressurized air. The glass slide was treated with a UV ozone cleaner at 100 °C for 30 min. Two chambers (SecureSeal™ hybridization chambers, Grace Bio-Labs, USA) were attached to the slide and heated at 100 °C for 60 s. Each chamber was first washed with 1 M KOH for 60 min at room temperature and rinsed three times with $H_2O$ and three times with PBS buffer. BSA-Biotin (1 mg/ml) was added to the chamber. After 20 min of incubation, the chamber was washed four times with PBS buffer to remove the excess of BSA-biotin. Then, the chamber was incubated with Neutravidin (1 mg/ml in PBS) for 10 min. Finally, the chamber was washed three times with PBS buffer and three times with NEB3-Tx-Tq-PCA buffer before adding the sample.

For the measurements, 5 pM of the annealed oligonucleotides (see Table S1) were dissolved in the NEB3-Tx-Tq-PCA buffer that was previously mixed with the 50× PCD buffer in a ratio of 50:1. After waiting for 5 min to immobilize the sample, the chamber was rinsed. Then, the sample was illuminated in alternating laser excitation (ALEX) with the green excitation exposing the sample for 100 ms, followed by the red laser again exposing for 100 ms. This pattern was repeated for 100 s.

#### 1.7 Data Analysis

##### 1.7.1 Lifetime Calculation

The data obtained with the TCSPC module was analyzed using a custom-made software built in Python, called smPyFLIM. The software counts with two modules, one to analyze time traces, and one to open and process FLIM images. The scripts needed to run the respective graphical user interfaces (GUI) can be found in https://github.com/alanszalai/smPyFLIM. The module that analyzes time traces reads .ptu, .fifo and .csv files, while the FLIM module opens .ptu files recorded with SymPhoTime64 (PicoQuant GmbH, Germany). The home-built software includes self-written functions, and built-in functions available from public libraries (such as NumPy, Matplotlib, or SciPy, as well as functions provided online by PicoQuant GmbH, Germany). In all the traces analyzed in this work, intensity and/or temporal thresholds were used to exclude photons arriving from the sample during time lapses where the fluorophore was in a dark state (*e.g.* in between blinking events, or after dye photobleaching). To obtain fluorescence lifetime traces as those shown in Fig. 1c, we set a temporal bin to sequentially gather photons that arrived during the acquisition. Furthermore, traces of the distance z to graphene and the bending angle θ were obtained from the fluorescence lifetime traces by using Eqn. 1 and 2 from the manuscript, respectively. The parameters used in these equations can be found in Table S2. The time binning was determined for each analyzed sample looking for the best spatio-temporal resolution balance. In the traces used for Fig. 1c and 1d, we aimed to only study the spatial resolution, and therefore we adapted the time bins to reach, on average, 1000 photons per time window. The time bin range used for these traces was ~30-150 ms. In the traces from enzyme-related experiments, we used a time binning of 50 ms, in order to detect dynamic bending or translocation events without compromising the spatial resolution. Here, the minimum number of photons per bin was around 250-300, while on average we obtained ~800-1000 photons per time bin.

In the lifetime calculation, we accounted for the contribution of the IRF (instrument response function) by performing a reconvolution analysis, which is also included in smPyFLIM. The IRF was acquired by detecting the reflection of the laser on a glass coverslip. Then, the convolution of the IRF with a monoexponential function with a background was used to fit the experimental data using a least-squares algorithm. Over weeks, the IRFs remained stable. However, we obtained IRFs routinely to minimize fitting uncertainties.

##### 1.7.2 Axial Localization Precision

The height histograms from Fig. 1e and 1h, as well as those used to obtain the parameters plotted in Fig. 1d, were fitted using single-peaked Gaussian functions, since there was no evidence of multiple populations resolvable by GETvNA. The widths of the distributions within single-molecule traces (Fig. 1c and 1d) were smaller than the widths obtained gathering data from many molecules and samples (Fig. 1e and 1h), as discussed in the main text and in section 2.1. The latter widths were typically between 8.5 and 10.5 Å. Therefore, we used 8.5 Å as the minimal width for stable populations in the histograms from Fig. 2b, 2d, 3e, and 3f, where hundreds of molecules from different areas and samples were analyzed together. In the histograms obtained at the single-molecule level, like those from Fig. 2b, 2c, 2d, and 3i, smaller boundaries were used for the distribution width, namely the maximum precision achievable, given by the Cramér–Rao bound (CRB). To calculate the CRB, we used the formula deduced by Thiele *et al.*[6] for the minimum theoretical error in a single-molecule lifetime measurement, and propagated it using $\sigma_{z,CRB} = \left|\frac{\delta z}{\delta \tau}\right| \times \sigma_{\tau,CRB}$, where $\sigma_{\tau,CRB}$ is the maximum precision obtained with the formula by Thiele et al. and $\left|\frac{\delta z}{\delta \tau}\right|$ is the derivative of $z$ with respect to $\tau$, which can be obtained by deriving Eqn. 2. This propagation assumes that $\tau_0$ and $d_0$ (that can be measured independently in separate experiments) have no uncertainty, and that the main source of error originates from the single-molecule lifetime measurement. This approach to calculate the CRB for $z$ through GET measurements was already done in a previous study from our lab[7]. $\sigma_{z,CRB}$ depends on $N$ (number of photons) and $SBR$ (signal-to-background ratio). Since the energy transfer from single molecules to graphene modulates the $SBR$ (the closer to graphene, the smaller the $SBR$), to plot the curve from Fig. 1d we obtained $\sigma_{z,CRB}$ for fixed $N = 1000$, $\tau_0 = 3.51\, ns$, as well as $SBR_{z=\infty} = 75$, and obtained $SBR$(z) through $SBR(z) = SBR_{z=\infty} \times \frac{1}{\left(\frac{d}{d_0}\right)^4 + 1}$. In Fig. S6, the CRB $\sigma_{z,CRB}$ is depicted as a function of the distance z of a fluorescent dye and a quencher for $SBR_{z=\infty} = 10$ as well as $SBR_{z=\infty} = 75$, and the previously mentioned parameters.

##### 1.7.3 Bulge and A-tract Angles

As laid out in the main text, a bending angle $\theta$ can be derived from a measured distance z to graphene by using Eqn. 2. Whereas the same experimentally determined value L is applied for all our measurements, the distance of the kink depends on the particular measurement.

$z_{bulge}$ is inferred from the experimentally determined length of the 37 bp construct (37mer + 73mer) as depicted in Fig. 1h, minus the expected length of the linker plus dye molecule, the latter being estimated from a comparison between external and internal labeling. More precisely, the experimental height of two constructs, namely a 45 bp construct (45mer ATTO 542 + 136mer) externally labeled at the last base pair, and a 66 bp construct (66mer internal ATTO 542 + 136mer) internally labeled at the 45th base pair, was determined. In the case of external labeling, the linker was assumed to protrude rigidly outside the DNA, as a virtual extension of its backbone; this is justified by ATTO 542 being negatively charged, and hence repelled by dsDNA. In the case of internal labeling, the linker protrudes perpendicularly to the vertically oriented dsDNA segment (see Fig. S10) due to electrostatic repulsions, therefore contributing only marginally to the measured height. Then, the average angle between the $z$-axis and the last base pairs of the 45 bp construct was inferred from MD simulations, as explained also in Fig. S10. Considering the difference in height between the two constructs (1.1 nm) and the obtained angle, it was possible to calculate the expected full length of the linker plus dye molecule: 1.2 nm. This value is in agreement with earlier reports by Hillisch *et al.*[8]. The same angle was estimated analogously for the upper segment of the 36 bp construct, as reported in Fig. S8, and from that the expected length of the z-projection of the linker plus dye for the 37 bp construct was calculated, assuming the angular orientation will not differ significantly from the one of the 36 bp system. In the end, the estimated height of the bulge corrected for the linker length was: $z_{bulge} = 14.5 - 1.1$ nm = 13.4 nm.

For the A-tract, which has its center at a height of 27 bp from graphene, $z_{A-tract} = \frac{27}{28} * z_{28bp} = 10.4$ nm is used instead of $z_{bulge}$ (see Fig. S14). For the lower and upper boundaries of the A-tract (used in Fig. 4d-f and Fig. S26-28), we used $\frac{24}{27} * z_{A-tract} = 9.2$ nm and $\frac{30}{27} * z_{A-tract} = 11.5$ nm, respectively. This approximation is justified as between 24 bp and 30 bp the respective dsDNA segment will only be slightly inclined and, hence, is assumed to be straight.

For the experiments involving Endo IV, the applied way for calculating bending angles is introduced in the following section.

##### 1.7.4 Enzyme Measurements

In the absence of Endo IV, a stable state with a monoexponential fluorescent decay was measured, exhibiting a single fluorescent lifetime, as it is visualized in Fig. 3b. The addition of the enzyme resulted in the observation of more than one state in the respective fluorescence intensity traces. Here, a window was set in the fluorescent intensity trace around the intensity of each visible state. The fluorescent lifetime corresponding to the state was obtained by fitting the fluorescent decay (see section 1.7.1) of all the photons within this window (see Fig. S20). Besides the assignment of common states in intensity space, states differing in their fluorescent lifetimes by less than ± 0.1 ns were summarized as the same state within one trace. From the traces exhibiting a switching between states based on the obtained fluorescence lifetimes, the smallest and the largest lifetimes were selected, converted to distances z to graphene by Eqn. 1 and depicted in the respective histograms. Here, the smallest and the largest fluorescence lifetimes are referenced by the lower and upper state, respectively. For all angle calculations, the respective distance $z_{AP} = 15.1$ nm of the AP site as given by the measurement of the 66mer (annealed to the 136mer) with an internal ATTO 542-labeling at nucleotide 45 (see Table S1), which is the exact position of the AP site, is used (see Fig. S10c).

40% of the analyzed dye molecules with AP site and PTO modification showed a dynamic switching between the states. From those constructs exhibiting dynamic changes among states, 71% showed two states whereas for 29% there were three visible states considering their fluorescent lifetimes (*cf.* Fig. 3c, d). In Fig. S20 and S22, we present additional examples with intensity-time traces, the applied thresholds, and the respective fluorescence decays.

Our measurements suggest that for the three-states case, there is an unbent, a prebent, and a fully bent state.[9] We come to this conclusion since the angular value of the intermediate state is always closer to the state with the largest angular value as well as that transitions to this intermediate state occur more likely to and from the, supposedly, fully bent state.

For data of the AGT enzyme, the fluorescent lifetime was determined for each 50 ms time bin. Subsequently, the respective distance z to graphene was derived by the use of Eqn. 1. The histogram of the step sizes was determined after filtering as it is outlined in section 2.3.

#### 1.8 Molecular Dynamics Simulations

All molecular dynamics (MD) simulations were performed using NAMD2.14[10], the bsc1parameter set for DNA[11], tip3p water model[12], and a set of CUFIX corrections to ion-nucleic acid interactions[13]. Graphene atoms were modeled as “ca” atoms from the gaff2 force field[14]. Multiple time stepping was used[15]: local interactions were computed every 2 fs, whereas long-range interactions were computed every 6 fs. All short-range nonbonded interactions were cut off starting at 1 nm and completely cut off by 1.2 nm. Long-range electrostatic interactions were evaluated using the particle-mesh Ewald method[16] computed over a 0.1 nm spaced grid and an interpolation order of 4. Switching functions to the non-bond interactions within the cutoff distance were turned “off”. The “scaled1-4” non-bonded exclusion policy was used with a scaling factor of 0.833. SETTLE and RATTLE algorithms were applied to constrain covalent bonds to hydrogen in water and in non-water molecules, respectively[17,18]. The temperature was maintained at 295 K using a Lowe-Andersen thermostat with a cutoff radius of 2.78 Å for collisions and the rate of collisions at 50 $ps^{-1}$. Constant pressure simulations employed a Nose-Hoover Langevin piston with a period and decay of 400 and 200 fs, respectively[19]. Energy minimization was carried out using the conjugate gradients method[20]. Atomic coordinates were recorded every 9.6 picoseconds, unless specified otherwise. Visualization and analysis were performed using VMD[21] and MDAnalysis[22].

A graphene sheet consisting of 3 layers was generated using VMD’s nanotube plugin[21]. The DNA molecules were built using the mrDNA package[23]. Each DNA construct contained a 5-nucleotide ssDNA fragment at the bottom. In the starting configurations, the dsDNA part of the constructs was placed perpendicular to and in close proximity with the graphene sheet. The ssDNA fragment was suitably rotated to avoid any contact with the sheet, keeping the dsDNA perpendicular to the sheet. In total three systems were simulated, distinguished by the size of the DNA construct. Each construct contained a 5-nt ssDNA overhang connected to a dsDNA duplex of 31, 51, or 66 base pairs. The DNA sequences of the duplexes were the same as in the corresponding experiments. The DNA constructs were submerged in a 50 mol $kg^{-1}$ NaCl solution.

Throughout all simulations, the graphene atoms were harmonically restrained to their initial coordinate, the spring constant of each restraint being 1 kcal $mol^{-1}$ $Å^{-2}$. After 4,800 steps of energy minimization, the system was simulated in the constant pressure and temperature ensemble and in the presence of an elastic network of restraints acting on the double stranded part of the DNA construct.[24] This allowed the ssDNA to form contacts with the graphene sheet. Subsequently, both position restraints and elastic network bonds on the DNA were completely removed and the system simulated in the constant pressure and temperature ensemble for a microsecond.

1.9 Comparison with Crystal Structure in Endo IV Experiments

To calculate the DNA bending angle, the helical axes of the two segments of the DNA separated by the AP site were determined. The 3DNA web package (http://web.x3dna.org/)[25] was used to find the origin of the base reference frame for each base pair. For each segment, we used the origins as a reference to calculate the distance to the rest of the origins and a line was fitted to the X, Y, Z coordinates using this distance as the independent variable. The axis is determined by the [X, Y, Z] coordinates obtained in the fit. Finally, to estimate the angle between the two helices, a normalized vector parallel to each helix was computed and the respective angle was calculated as the arccosine of the dot product of these vectors.

The PDB structure 2NQJ[3] was used to calculate the DNA bending angle created by Endo IV, *cf*. Fig. S24. To estimate the intrinsic bending of the DNA at an AP site, the averaged structures determined by NMR[26] for the alpha (PDB 2HSL) and beta (PDB 2HSS) anomers were used, *cf*. Fig. S19.

### 2 Supplementary Text

#### 2.1 Single-Molecule vs. Ensemble Precision

When comparing Fig. 1d and Fig. 1e, the reader will note a significant difference in the obtained axial precision. This leads to two questions: i) Why is the precision/uncertainty of the distribution in Fig. 1e substantially larger than the one in the respective case in Fig. 1d? ii) While Fig. 1d shows that the precision worsens for longer dsDNA segments, Fig. 1e depicts the contrary, with an improving precision for longer dsDNA segments.

The discrepancy underlying i) can be explained by the fact that in a single trace exhibiting one stable state the photon count is solely shot-noise limited, while the histogram of an ensemble of single molecules as in Fig. 1e accounts for the local variations in graphene quality. This heterogeneity comprises, *e.g.* holes and defects in the crystal structure, which can hinder the quenching properties of graphene, thereby reducing the effective $d_0$, as well as small bilayer areas. Each result could be reliably reproduced with independent graphene samples, showing that these heterogeneities do not significantly affect the final population-averaged result. Though, it is reasonable to assume that they can increase the variability between different analyzed single molecules. For ii), two scenarios must be distinguished: the resulting fluorescence lifetime from a short-noise limited trace is estimated more accurately, *i.e.* the respective uncertainty is smaller, the smaller the respective lifetime value. The uncertainty in lifetime propagates for a photon count of 1000 and a lifetime of 3.0 ns almost linearly with the distance z. For this, shorter dsDNA segments mean higher quenching of the attached fluorophores, lower fluorescence lifetimes and, consequently, a smaller uncertainty on the resulting distance z. In contrast, when a population of single molecules is considered, one has to take into account that fluorophores attached to shorter dsDNA segments, and hence closer to graphene, sense a comparatively smaller area of the underlying graphene sheet. Therefore, the effective $d_0$-value for this smaller area reflects more strongly local variations of the graphene sheet leading to an increased uncertainty compared to longer dsDNA segments. More precisely, the area of quenching $A_q$ of the graphene sheet sensed by a molecule scales quadratically with its distance $z$ from the surface, as shown in the following formula: $A_q = \pi\left(\sqrt{\frac{1}{1-\beta}}-1\right)z^2 = \pi\, r_q^2$. Here, β represents the part of the total graphene area contributing to the quenching. For reference, a dye located at 14.0 nm above the surface (as for the 36 bp dsDNA segment) will receive 99% (*i.e.* β = 0.99) of the total quenching effect from an area of 5,542 nm$^2$. In contrast, with z = 19.3 nm (as for the 66 bp dsDNA segment) this value is 10,532 $nm^2$. This screening effect of a defined area of graphene is visualized in Fig. S7.

#### 2.2 Comparison of Accuracy Obtained by GET and FRET

As outlined in sections 1.7.3 and 1.7.4, a measurement of a distance $z$ is converted into a bending angle $\theta$ using a purely one-dimensional model for dsDNA. The two dsDNA segments separated by the kink can be modeled as two rigid cylinders allowing each cylinder to have an independent torsion angle (ϕ and ψ respectively) with respect to its axis, as well as a relative displacement of the two cylinders in the three directions ($x_0, y_0$ and $z_0$). In this case, the whole construct has six degrees of freedom, as depicted in Fig. S11.

In order to estimate $\theta$ with FRET, both dsDNA segments have to be labeled with dye molecules. Thus, the bending angle $\theta$ is estimated from the measured distance between acceptor and donor dye molecule. This distance depends on each one of the aforementioned degrees of freedom, and hence parallel experiments are required in order to estimate their values, increasing the complexity of both the measurements and the model.

In order to estimate $\theta$ with graphene energy transfer (GET), instead, only the upper dsDNA segment has to be labeled. Thanks to the planar geometry of graphene, its distance from the dye depends only on two degrees of freedom ($z_0$ and ψ - the torsion angle of the upper dsDNA segment). Additionally, given that the typical donor-acceptor distance is far greater in GET than in FRET, every variation in the distance between them due to the uncertainty in the additional degrees of freedom has a smaller relative effect. This is summarized in Fig. S11, where one can see that the inaccuracy introduced by using a one-dimensional model for dsDNA is roughly ten times bigger in the case of FRET compared to GET. The small value of the inaccuracy for GET, of the same order of magnitude as the experimental uncertainty, justifies the use of a one-dimensional model, allowing to estimate bending angles with single measurements on a single molecule, which is crucial, especially, when measuring dynamics (*e.g.* in Endo IV experiments).

#### 2.3 Translocation of AGT - Filtering of Steps

To quantify the steps of AGT monomers and clusters diffusing along dsDNA we performed a filtering step as sketched in Fig. 4g. This filter allowed us to consider only initial and final states (before and after the protein step) with heights that can be accurately measured. Remarkably, the CRB for ATTO 647N at a height of 9 nm (the average height of AGT measurements) and 1000 photons is 0.8 Å. Therefore, under those conditions, it should be possible to resolve movements of c.a. 3.1 Å, *i.e.* the z-projection of the inter-base pair distance for vertically standing dsDNA at z = 9 nm, considering a local angle of the main dsDNA axis with respect to the z axis in this region of around 20° (see dsDNA angles for the last six bp of a 36 bp system, *i.e.* z ~11 nm, in Fig. S8). However, this can only be achieved if the protein moves slow enough so that each state can be sampled with the required number of photons. Since both clusters and monomers showed slow and fast diffusive components, the spatio-temporal resolution achieved in our GETvNA measurements did not always allow for detecting single base-pair displacements. For example, in some fast events, as the one shown in the zoomed-in region from Fig. 4c, the protein has moved more than 5 nm (> 14 base pairs) in the axial direction within the duration of a single time bin (50 ms). In such scenarios, a fixed time bin of 50 ms impaired the possibilities of studying whether the protein made a stepwise movement or not. In other words, when the protein has moved too fast, the height corresponding to adjacent bins differs significantly, changes constantly, and therefore the measured position in each time bin only represents an average.

Strikingly, during the slow diffusion of the protein, it is possible to see displacements that correspond to < 1 nm (< 3 base pairs) jumps, as highlighted in Fig. 4c, and also visible in Fig. 4g. Thus, a temporal resolution of 50 ms demonstrated to be good enough to track the AGT translocation with single-base pair resolution in the slow mode. However, the optimal way to calculate the height associated with a protein step in the dsDNA axis direction must include a filtering step. First, a lower, spatial threshold was defined in order to detect a jump between different states. For this, we chose a value of at least 2.0 Å for $|z_{t_{i+1}} - z_{t_i}|$, *i.e.* for the absolute value of the difference in height between consecutive bins. This value lies above $2 * \sigma_{z,CRB}$ (N = 1000, $SBR_{z=\infty} = 75$, and $\tau_0$= *3.78 ns*), and it is still significantly below the value of a single base pair step. As a second step, a filter to select stable states needs to be applied. If the protein stays in a given state for only one time bin, it will contain some components of the previous and the subsequent state. On the other hand, if it stays for a long time, then the state will be sampled with multiple bins and its position can be well-estimated with the average position of all bins, except from the initial and last ones, which will have components of the previous and the subsequent state, respectively. For this reason, we chose stable states that lasted at least four bins, in order to have a minimum number of two points to determine the height of each state. Expectedly, only a small fraction of bins from every analyzed time trace were included in the analysis after this filter was applied, enabling us to obtain an accurate measure of the protein displacement. To increase the statistics, we allowed a temporal window between stable states having a maximum width of 10 bins, *i.e.* 0.5 s. This also enabled us to detect steps of 0 bp whenever the protein returned to the same base pair after visiting for a short period a neighboring one, as evidenced by the central peak of the histogram from Fig. 4h.

In Fig. S30, we show the effect of varying the threshold to detect jumps by AGT. The shown range for the lower, spatial threshold is limited to 1.8 Å - 2.2 Å, reflecting the values with a physical meaning, since smaller values overlap with $\sigma_{z,CRB}$ and would give rise to the detection of many false jumps. In turn, with a larger spatial threshold, the number of detected stable states lasting more than three bins drastically dropped. Additionally, larger values would decrease the probability of detecting jumps of one base pair, therefore distorting the shape of the ± 1 bp populations in the protein step histogram. In Fig. S30, the effect of varying the maximum width of the temporal window, is also shown. Here, it can be seen that a too large tolerance in the time window gap (> 5 s) starts distorting the positions of the peaks. On the other hand, the analysis gives similar results when varying the intensity threshold to detect jumps in the studied conditions.

In Fig. S31, we depict the protein step histogram for the data obtained with dsDNA lacking an A-tract.

#### 3 Supplementary Figures

**Fig. S1.**

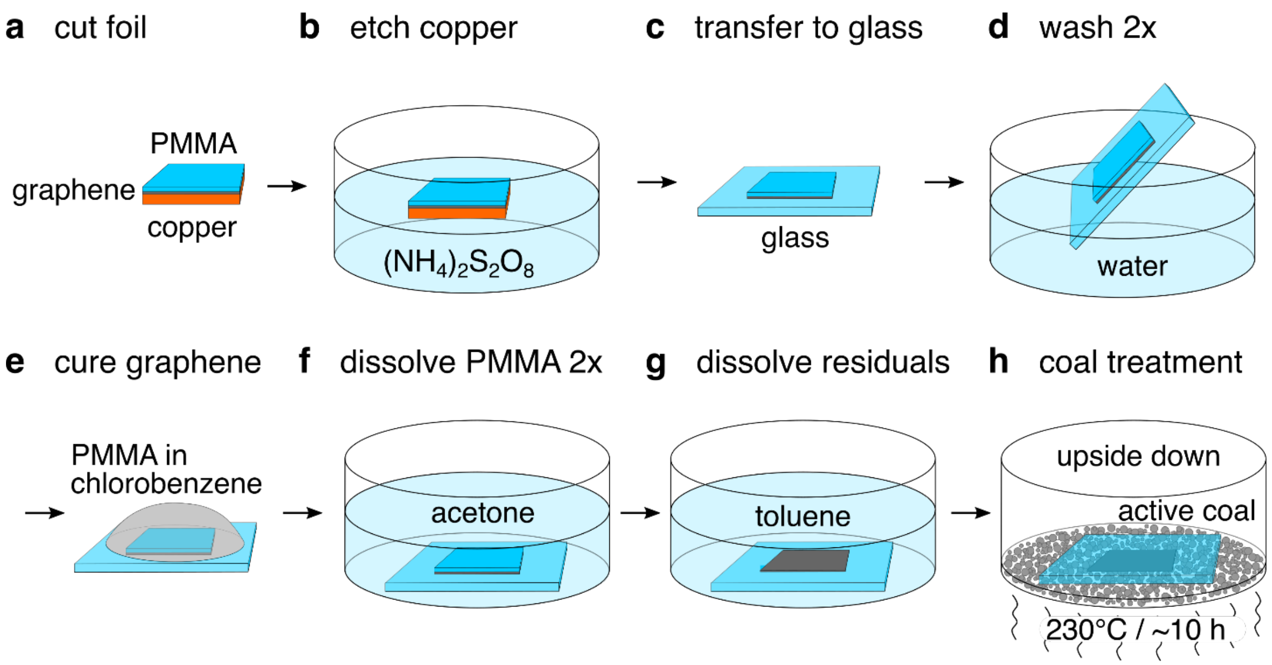

**Fig. S1. Workflow for graphene-on-glass transfer.** The sketches show the preparation of graphene-on-glass coverslips with the single steps denoted by the letters **a)** to **h)**. Adapted from Krause *et al.*[1].

**Fig. S2.**

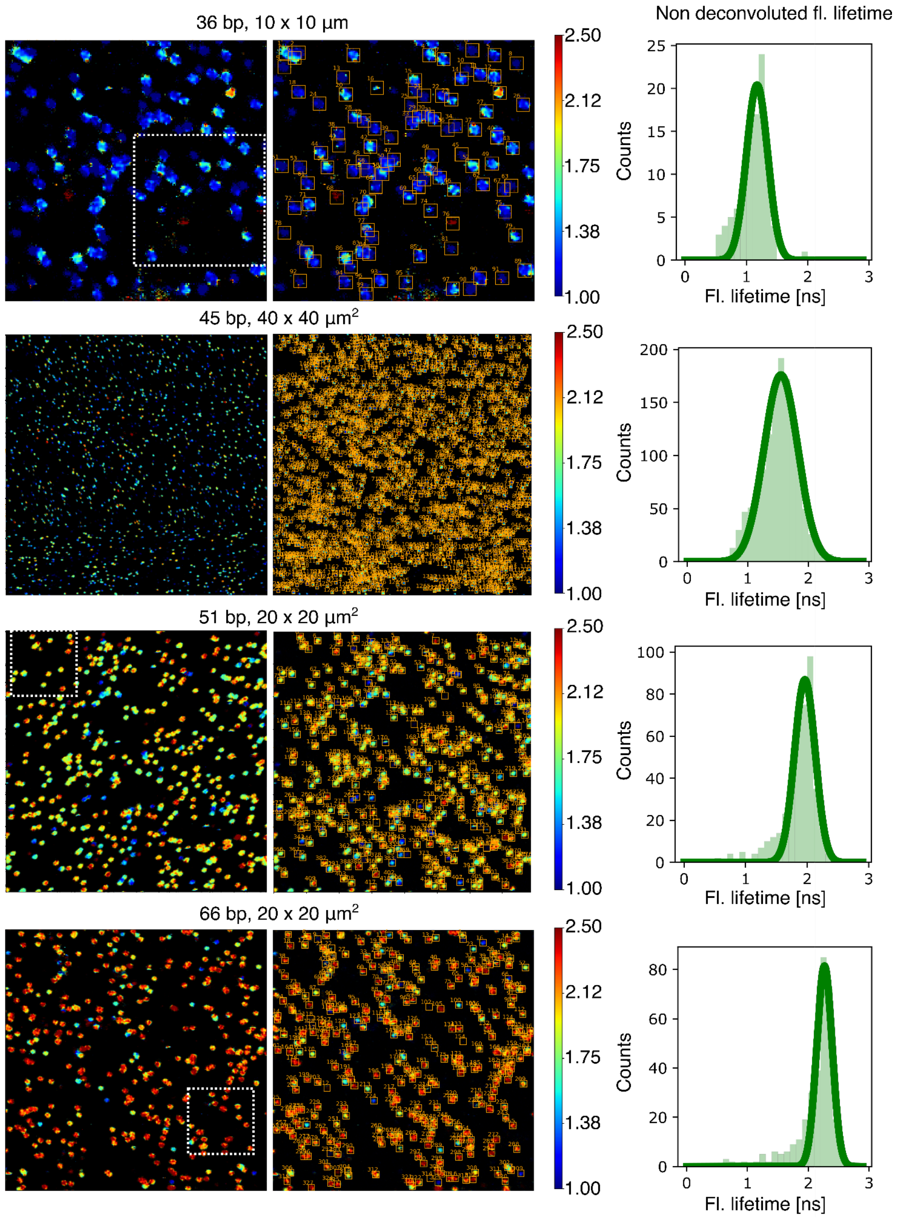

**Fig. S2. Extended field-of-views of hybrid DNA constructs immobilized on graphene.** From top to bottom: extended fields of view for systems containing dsDNA segments of 36 bp, 45 bp, 51 bp and 66 bp length. For the 36 bp, 51 bp and 66 bp cases the white dotted boxes mark the areas shown in Fig. 1b. For each case, the FLIM images are shown twice with the second image representing the results of an automatic peak detection algorithm. The non-deconvoluted fluorescence lifetime of each detected spot was computed, and the corresponding histograms are shown in the right panels. There, it can be seen that the fluorescence lifetimes are homogeneous for large areas. A shift to larger fluorescence lifetimes is observed when the length of the dsDNA segment is increased.

**Fig. S3.**

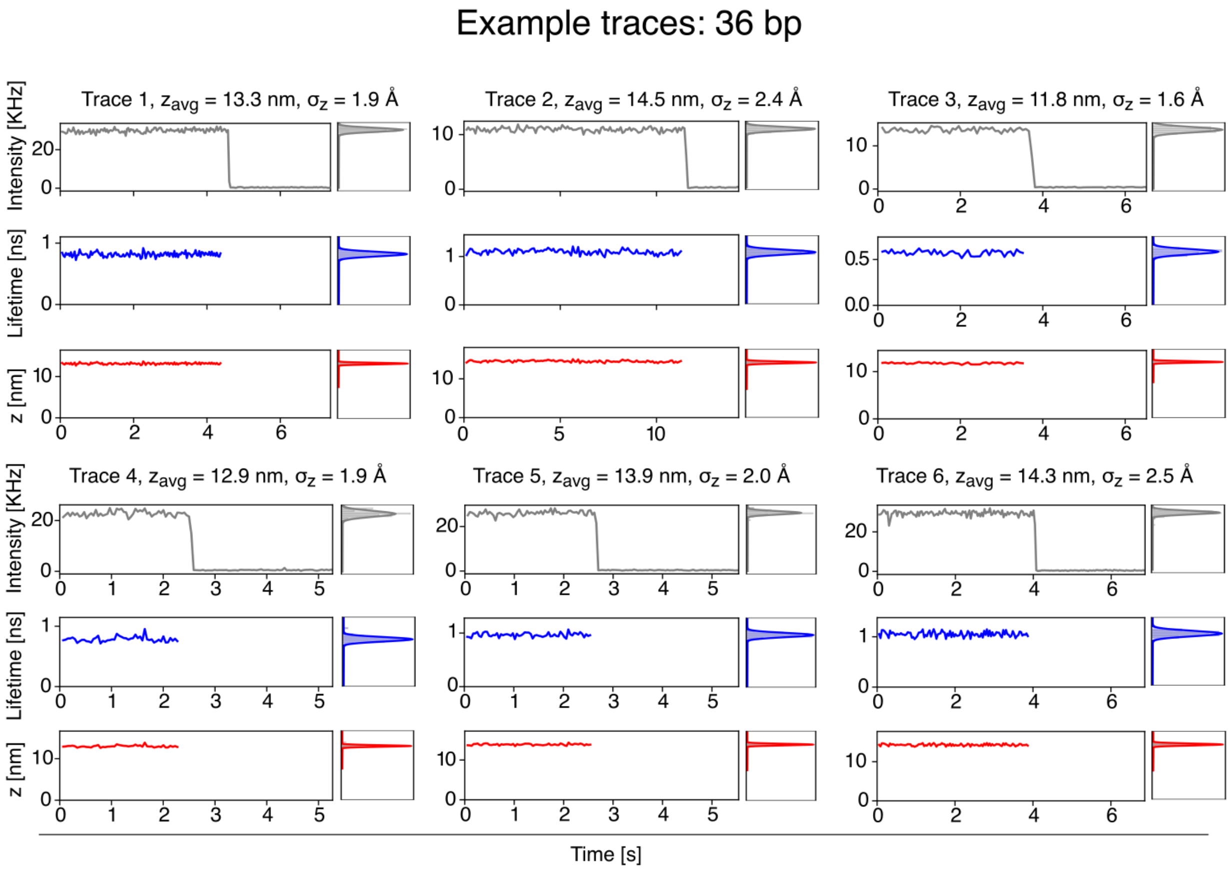

**Fig. S3. Exemplary time traces of systems containing a 36 bp-long dsDNA segment.** Six exemplary time traces are shown. For each case, the fluorescence intensity, the fluorescence lifetime, and the distance to graphene *z* are plotted. On the right of each plot, the corresponding histogram is shown. The average distance to graphene, as well as the axial localization precision, are presented on top of each trace.

**Fig. S4.**

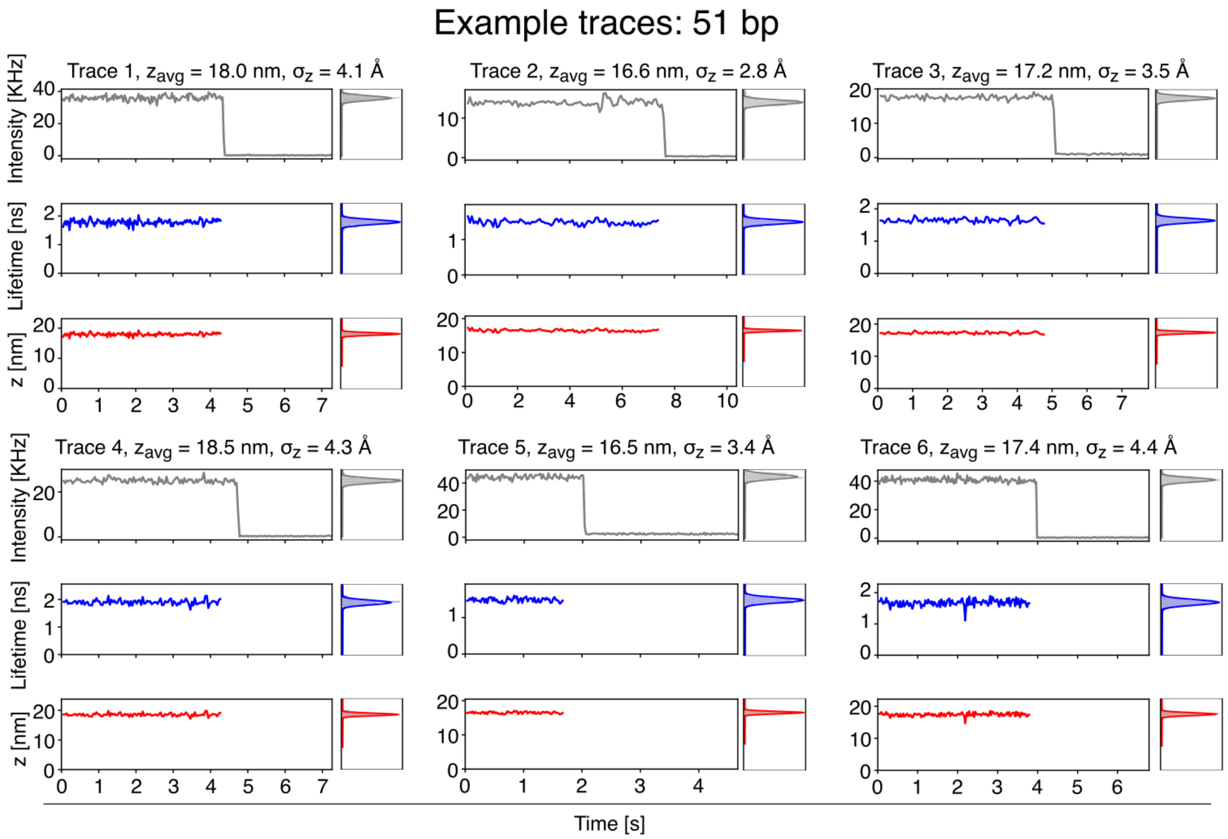

**Fig. S4. Exemplary time traces of systems containing a 51 bp-long dsDNA segment.** Six exemplary time traces are shown. For each case, the fluorescence intensity, the fluorescence lifetime, and the distance to graphene *z* are plotted. On the right of each plot, the corresponding histogram is shown. The average distance to graphene, as well as the axial localization precision, are presented on top of each trace.

**Fig. S5.**

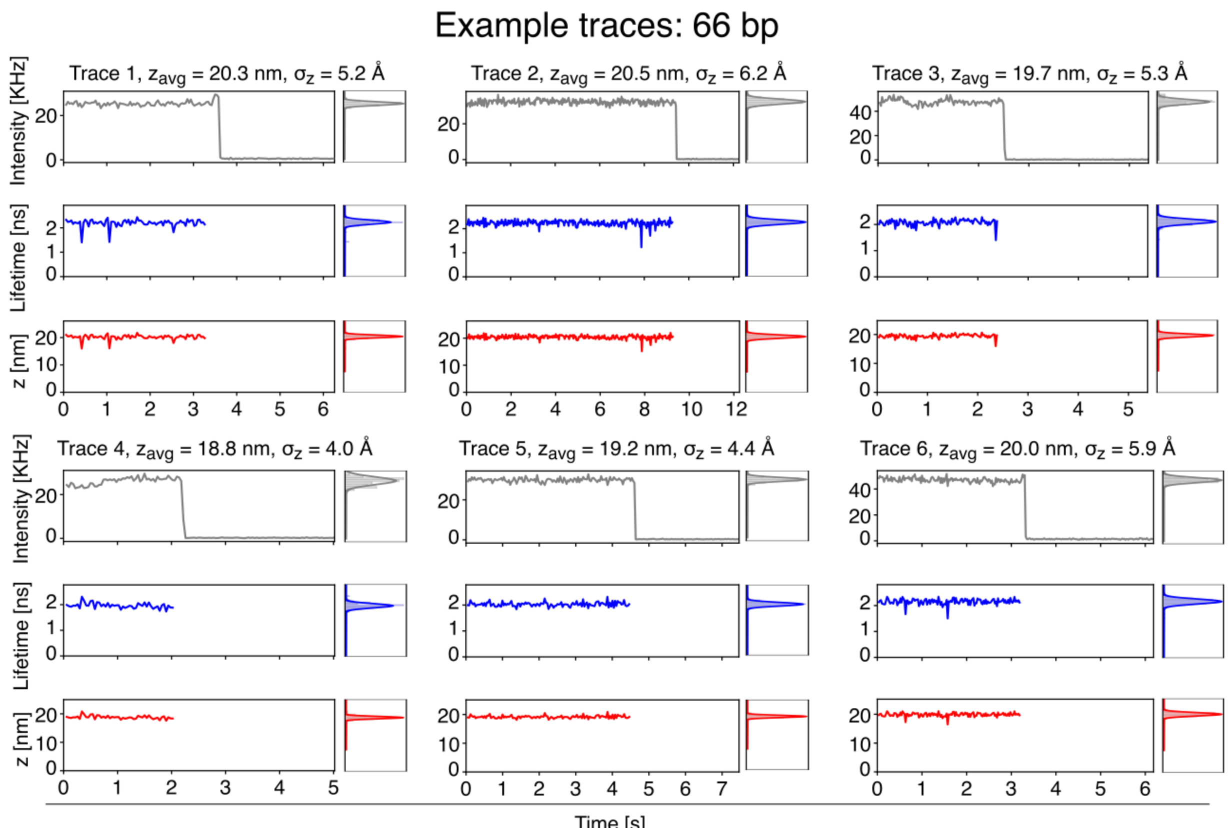

**Fig. S5. Exemplary time traces of systems containing a 66 bp-long dsDNA segment.** Six exemplary time traces are shown. For each case, the fluorescence intensity, the fluorescence lifetime, and the distance to graphene *z* are plotted. On the right of each plot, the corresponding histogram is shown. The average distance to graphene, as well as the axial localization precision, are presented on top of each trace.

**Fig. S6.**

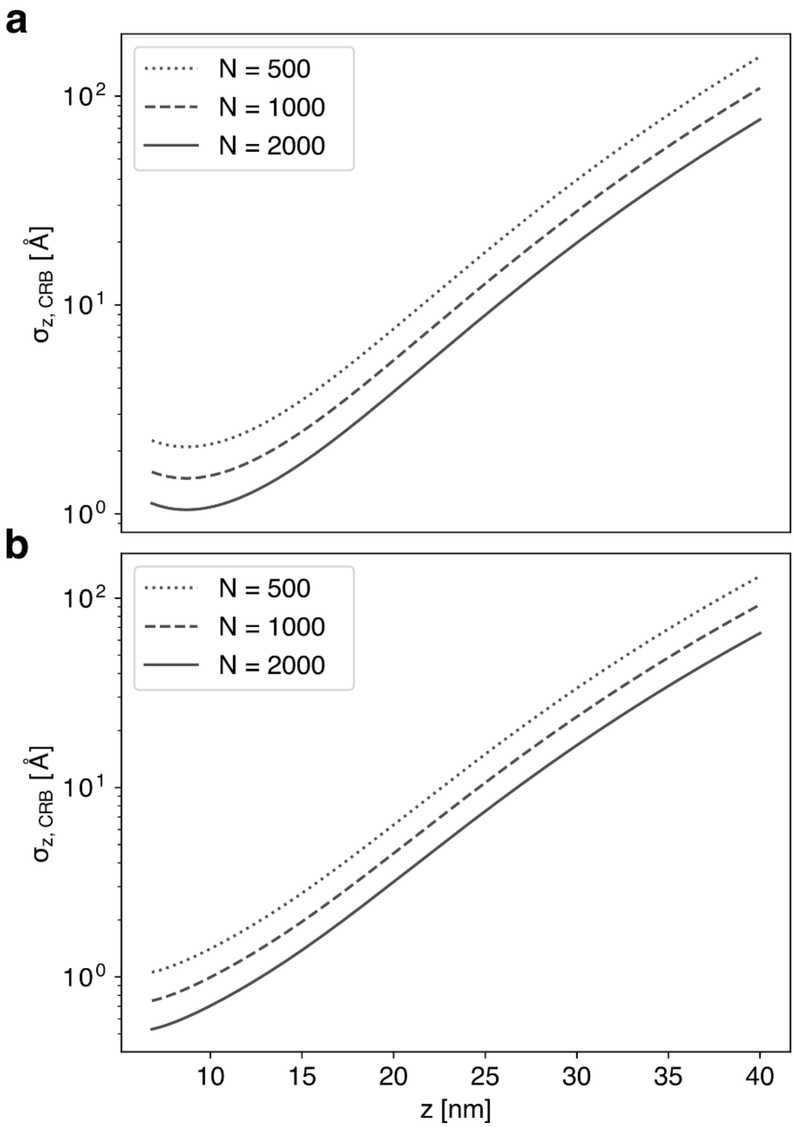

**Fig. S6. The axial precision $\sigma_{z,CRB}$ as a function of the distance to graphene z.** The dependency is shown for three different numbers of photons N. An unquenched fluorescence lifetime of 3.51 ns was used. **a)** $SBR_{z=\infty} = 10$ and **b)** $SBR_{z=\infty} = 75$.

Fig. S7.

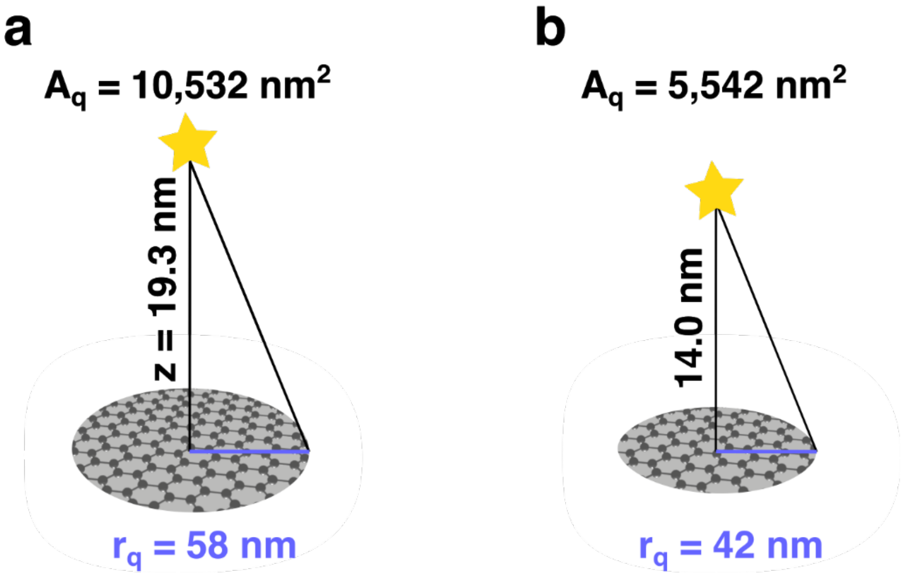

**Fig. S7. Comparison of the graphene area sensed by single molecules located at different distances z.** The sketches show how quenched fluorescent dyes positioned at larger distances to graphene are sensitive to larger graphene areas. In **a)**, $z = 19.3$ nm and in **b)** $z = 14.0$ nm. The respective quenching area $A_q$ as well as the radius $r_q$ of the corresponding circular area of graphene are depicted. Here, both quantities are related by $A_q = \pi\, r_q^2$. In both cases, $A_q$ represents the area of graphene responsible for 99% of the fluorescence quenching experienced by the dye as introduced in section 2.1.

**Fig. S8.**

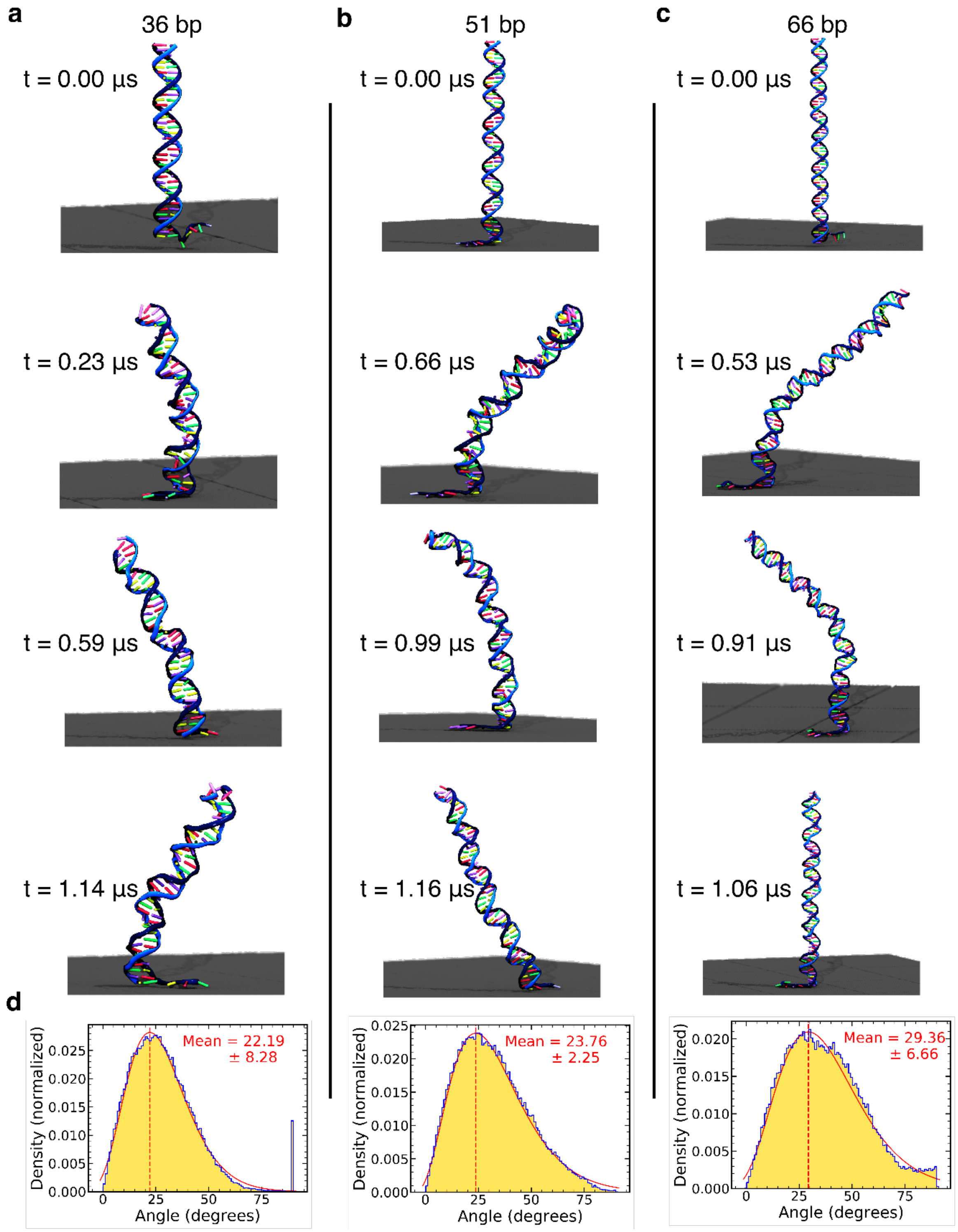

**Fig. S8. MD simulations of DNA constructs. a-c)** Snapshots illustrating MD simulations of the 31, 51 and 66 base pair DNA constructs at the indicated simulation times. For visual clarity, water and ions are not shown. The DNA backbone is shown as a ribbon (blue) and the bases as sticks colored by the base type. The graphene sheet is shown as a smooth surface (gray). **d)** Distribution of the angle that a DNA duplex at the end of each construct forms with the z axis. To compute the orientation of the duplex, coordinates of N1 atoms of purine and N3 atoms of pyrimidine bases were recorded every 9.6 ps for a six base pair fragment of the constructs located two base pairs away from the construct's end. For every frame of the MD trajectory, the direction was identified from the covariance matrix of the atomic coordinates, as the eigenvector of the largest eigenvalue. A gamma distribution (red) was fitted to the distributions to obtain the mean values (red dashed line).

**Fig. S9.**

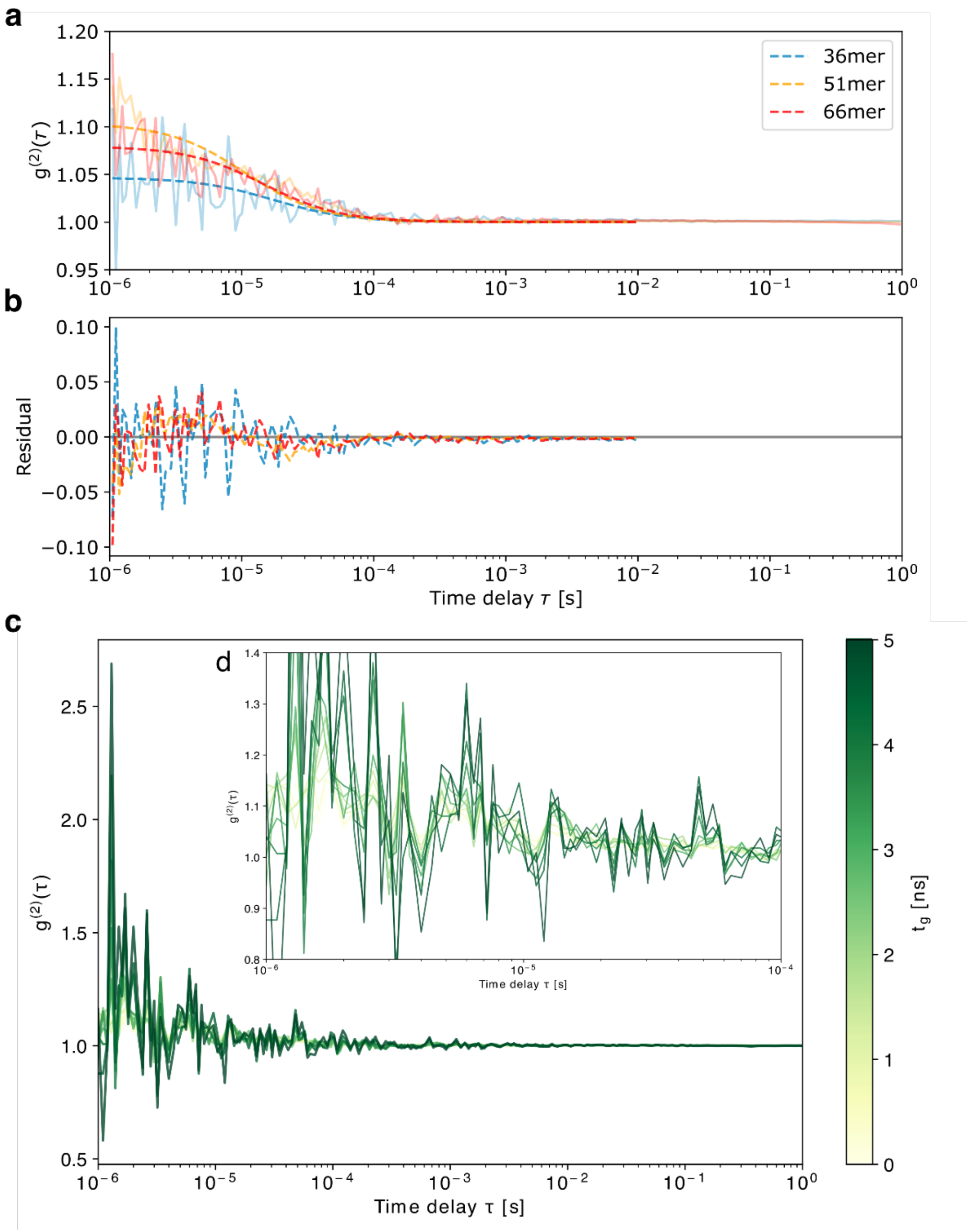

**Fig. S9. Autocorrelation of hybrid DNA constructs demonstrating its conformation.** In **a)**, each depicted autocorrelation function results from averaging 10 autocorrelated time-intensity traces. For clarity, the functions are fitted with a three-dimensional diffusion model as depicted by the dashed lines on top of each function. In **b)**, the respective residual is shown for each fit. **c)** visualizes the autocorrelation function of an intensity trace from one example of a 66mer showing a similar behavior as in a). Here, the microtimes were time-gated with different gating times $t_g$ to distinguish dynamical processes from photophysics.[27] The zoom-in in **d)** shows that the resulting amplitudes in a) can be associated with a dynamical process, whereas photophysics can be excluded.

**Fig. S10.**

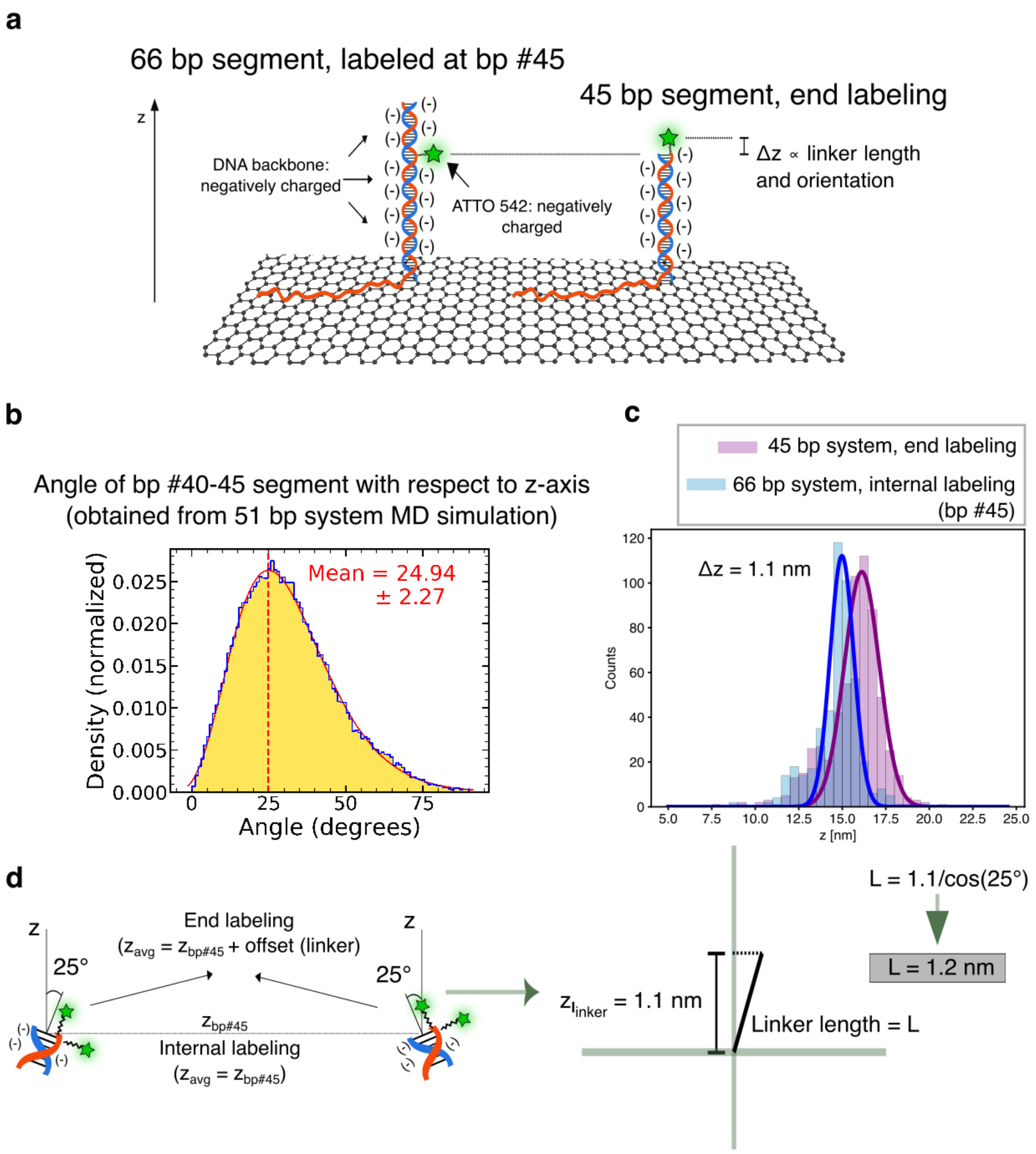

**Fig. S10. Determining the contribution of the linker to the axial position of single molecules. a)** Sketch showing the systems used to estimate the contribution of the linker. Left: system containing a dsDNA segment with 66 bp, internally labeled at base #45. Right: system containing a dsDNA segment with 45 bp, labeled at one of its end bases. The negative charges are highlighted, since they are responsible for extending outwards the negatively charged dye (ATTO 542), which is attached through a six-atom carbon linker. **b)** Distribution of angles with respect to the *z*-axis obtained for the 40-45 bp segment from the MD simulation trajectory of the 51 bp system. The methodology to calculate this angle was analogous to the one described in the caption of Fig. S8d. **c)** Height distributions for the two systems described in a). **d)** Representation of the trigonometric calculations performed to retrieve the linker length assuming a model where the linker is stretched, extending outwards of the dsDNA segment (following the direction of the dsDNA segment for the end-labeled case, and oriented perpendicularly for the internally labeled scenario). The 25° angle used was obtained from the histogram shown in b), and the 1.14 nm height difference was extracted from the histograms shown in c).

**Fig. S11.**

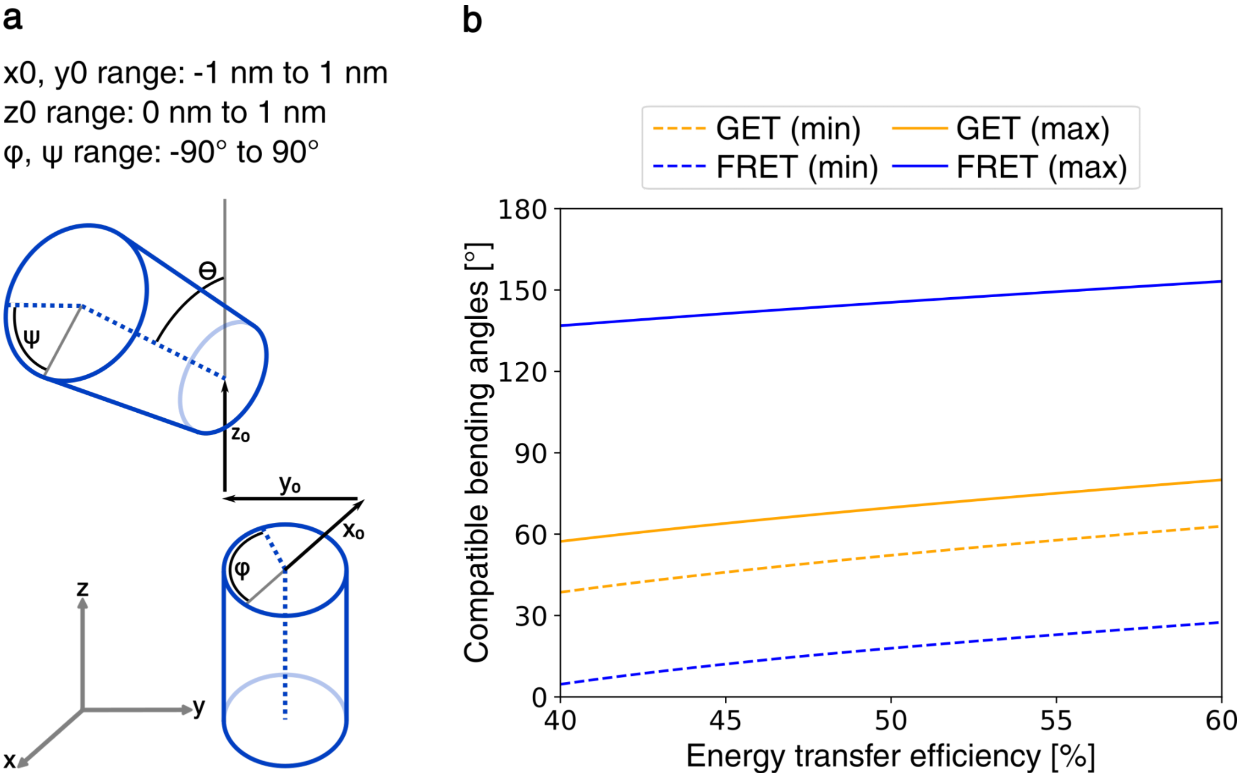

**Fig. S11. Comparison between ranges of bending angles compatible with a given measured energy transfer efficiency for GET and FRET. a)** Sketch showing a simple model for kinked dsDNA, consisting of two rigid cylinders which can rotate around their respective axes (with torsion angles ϕ and ψ, respectively). They move with respect to each other ($x_0, y_0$ and $z_0$ represent the displacements in three dimensions of the bottom of the upper cylinder with respect to the top of the lower one), and bend by an angle $\theta$. **b)** Plot showing the minimum and maximum bending angle $\theta$ compatible with a given energy transfer efficiency between 40% and 60%, for GET and FRET. The model shown in a) was considered for the calculations, with 0.34 nm base pair (bp) length, 1 nm dsDNA radius, and 10.5 bp per double helix full turn as physical parameters. Two different labeling strategies were chosen for the two methods: for GET, the kink was positioned at 36 bp distance from graphene, and the dye at 30 bp distance from the kink, in the upper segment; for FRET, the two dyes were both positioned at 8 bp distance from the kink. $d_0$ for GET and $r_0$ for FRET were set at 17.7 nm and 5 nm respectively. For each value of the energy transfer efficiency $x_0, y_0$ were varied from -1 nm to 1 nm in steps of 0.5 nm, $z_0$ was varied between 0 nm and 1 nm (with no steps in between), ϕ and ψ were varied from -90° to 90° in steps of 1°. For each combination of these parameters, the value of $\theta$ leading to the chosen energy transfer efficiency was computed. The plotted minimum and maximum values of $\theta$ refer to all the possible combinations of parameters.

**Fig. S12.**

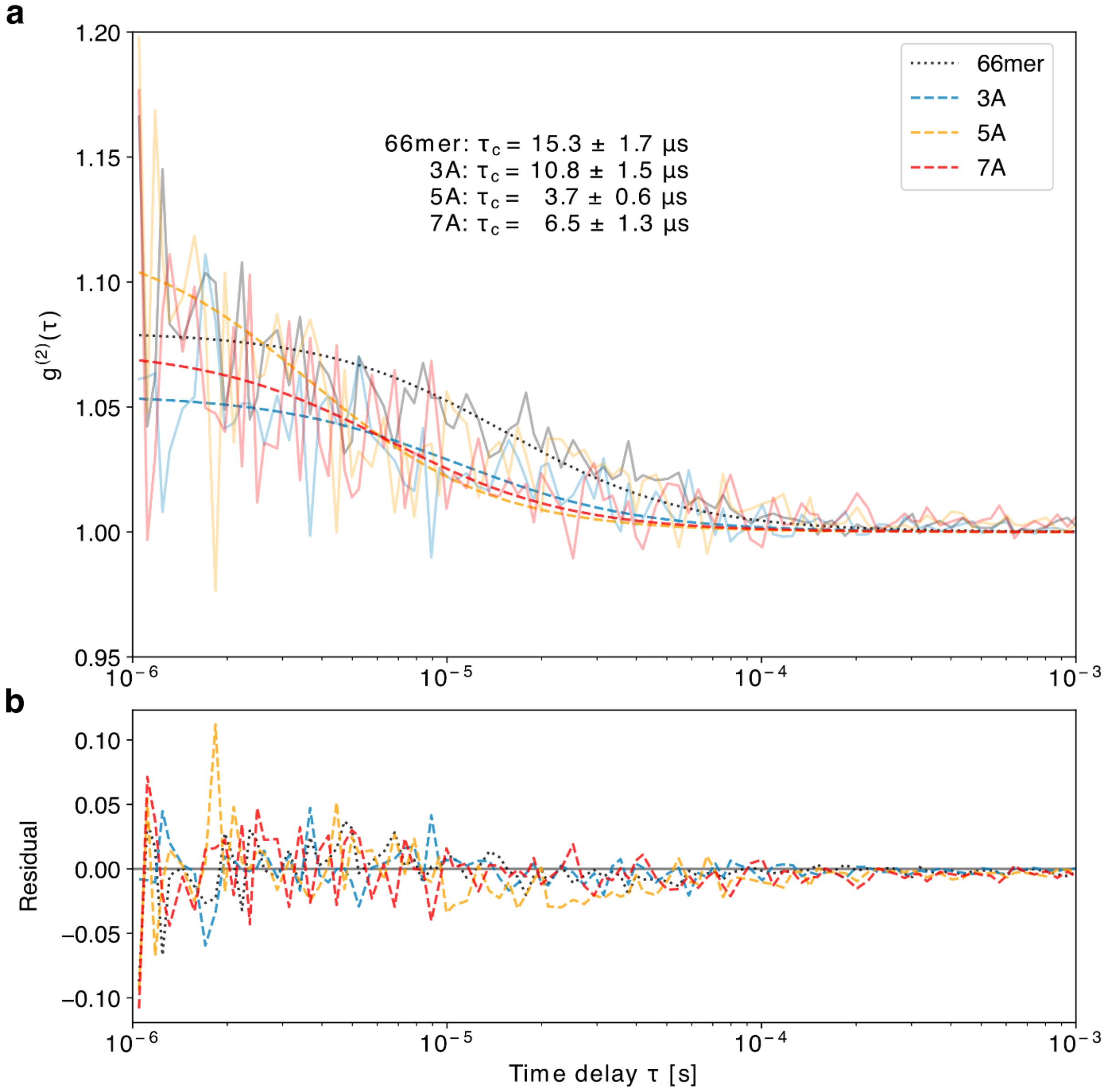

**Fig. S12. Autocorrelation of 3A, 5A and 7A bulge systems as well as for the 66mer. a)** Each depicted autocorrelation function results from averaging 10 autocorrelated time-intensity traces. For clarity, the functions are fitted with a three-dimensional diffusion model as depicted by the dashed lines on top of each function. The corresponding correlation times $\tau_c$ are enumerated for the respective fitting functions. **b)** The respective residual is shown for each fit. As demonstrated in Fig. S9, the shown correlation amplitudes are associated with a dynamic process. Also, based on the correlation amplitudes being below

1.3, we assume that each molecule from a corresponding bulge system does not change its state during the measurement.

**Fig. S13.**

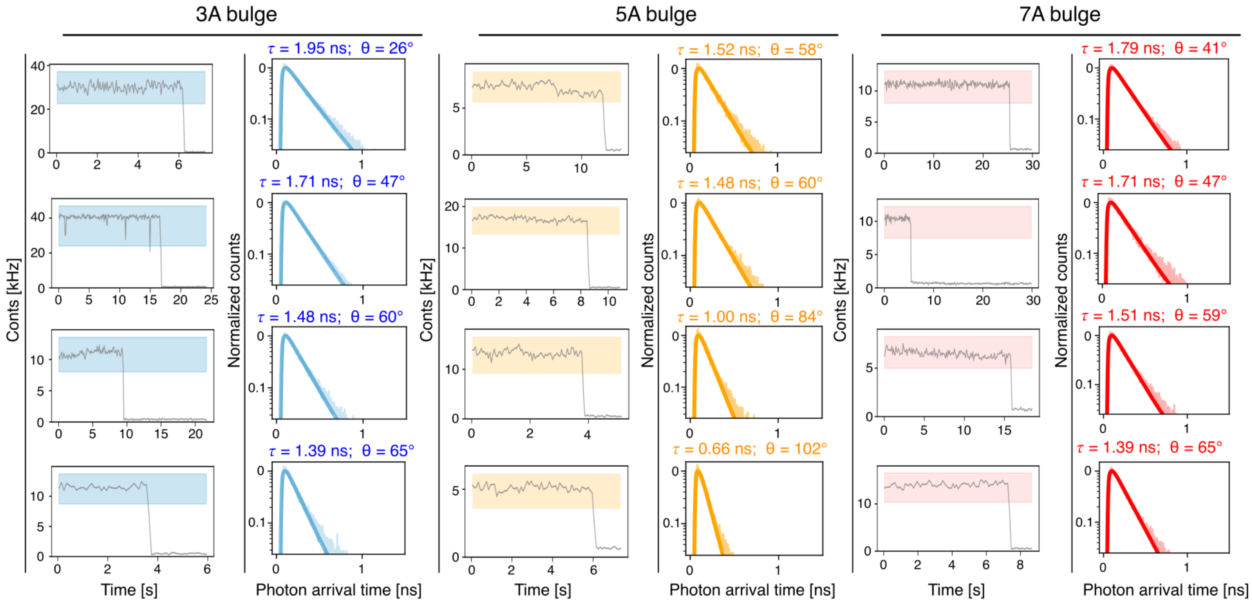

**Fig. S13. Exemplary time traces of systems where the dsDNA segment contained a bulge.** Four example time traces are shown for each bulge (3A, 5A, and 7A). The fluorescence intensity time traces are shown on the left, and the fluorescence decays and corresponding monoexponential fits on the right. For each case, the fitted fluorescence lifetime and the corresponding bending angle are shown on top of the fluorescence decay plots. The rectangles from the intensity time traces highlight the photons used to obtain the fluorescence lifetime decay curves.

**Fig. S14.**

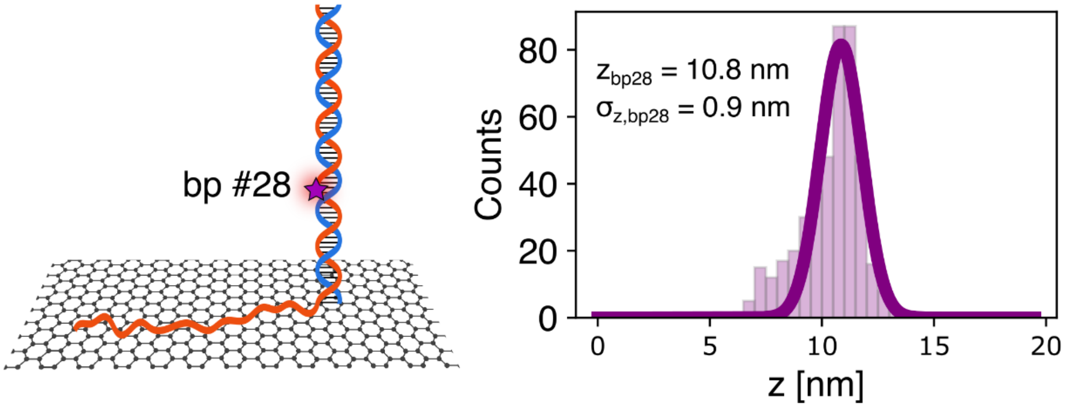

**Fig. S14. A-tract height determination.** Left: sketch of the internally labeled construct. The dsDNA segment was functionalized with ATTO 647N at one of the bases of bp #28. Right: height histogram for bp #28 obtained from 436 single-molecule traces.

**Fig. S15.**

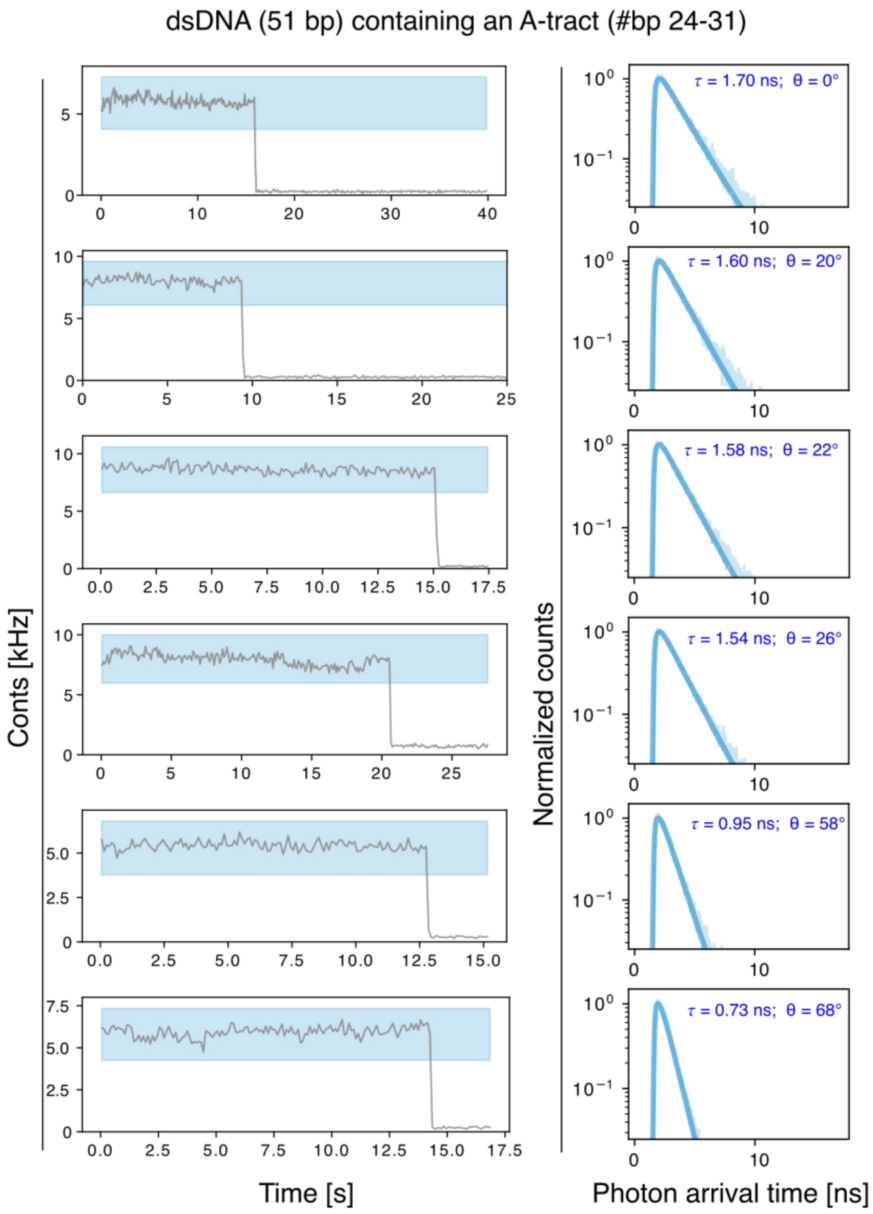

**Fig. S15. Exemplary time traces of systems where the dsDNA segment contained an A-tract.** The A-tract was located between bp #24 and #31 (counting from the dsDNA end facing graphene). Six example time traces are shown. The fluorescence intensity time traces are depicted on the left, and the fluorescence decays and corresponding monoexponential fits on the right. For each case, the fitted fluorescence lifetime and the corresponding bending angle are shown within the fluorescence decay plots. The rectangles from the intensity time traces highlight the photons used to obtain the fluorescence lifetime decay curves.

**Fig. S16.**

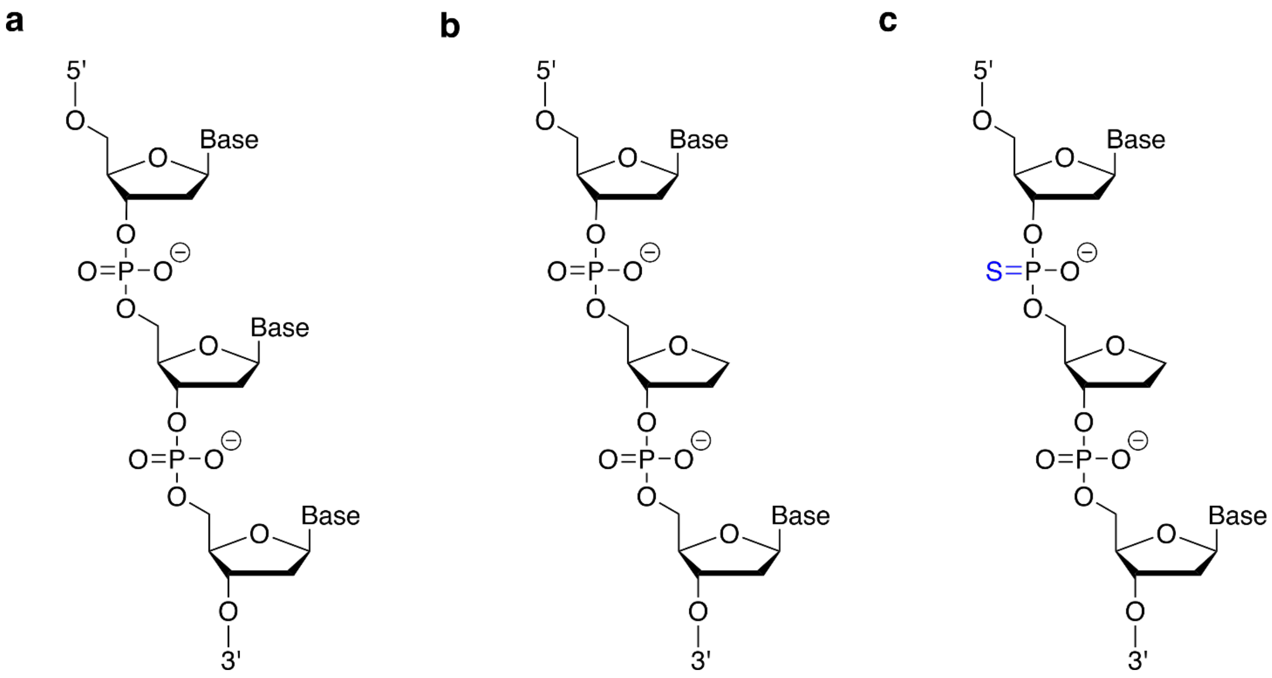

**Fig. S16. Chemical structures of DNA modifications. a)** Unmodified ssDNA. **b)** DNA strand with an AP site as used within this study. **c)** Additional PTO modification indicated in blue creating a DNA strand with AP site as well as a PTO modification.

**Fig. S17.**

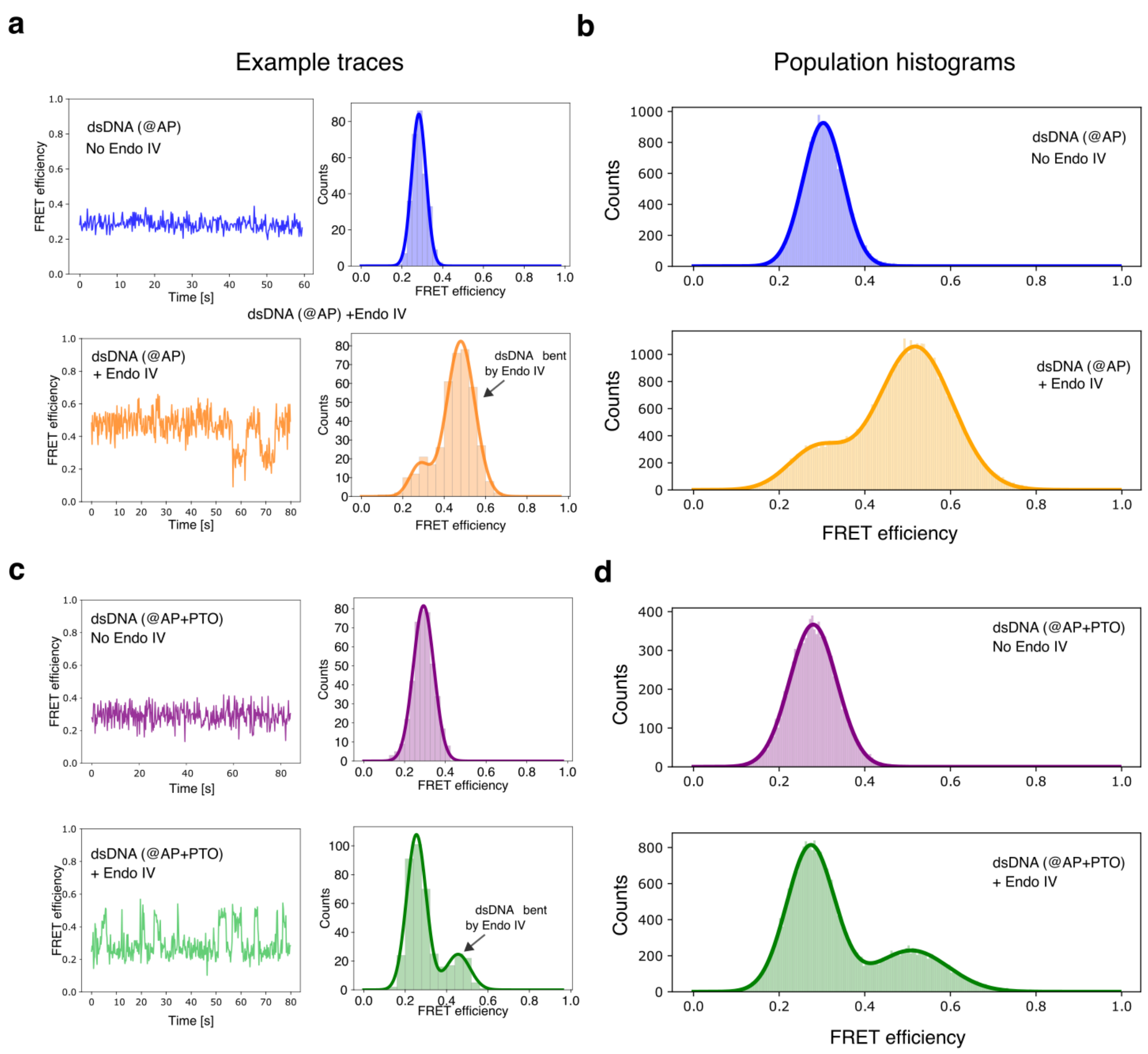

**Fig. S17. Intensity-based smFRET studies of dsDNA containing an AP site in the presence and absence of Endo IV.** The influence of having a PTO modification framed by the 5'-neighboring nucleotide and the AP site is evaluated. **a)** Example smFRET time traces of the system containing dsDNA with AP site without PTO modification, in the absence (top) and presence (bottom) of Endo IV. On the right, the histograms obtained from the shown traces are depicted. **b)** FRET efficiency histograms obtained from 75 (dsDNA containing AP, without PTO, in the absence of Endo IV) and 55 traces (dsDNA containing AP, without PTO, in the presence of Endo IV). Here, the FRET efficiencies obtained from every movie frame from all traces are computed together as independent

FRET efficiency values. **c)** and **d)** Same description as in a) and b), but for systems containing both AP site and PTO modification. 30 traces were analyzed for the population histogram without Endo IV and 66 traces for the case, where Endo IV was added to the solution. Due to the intensity-based measurement protocol, the histograms from b) and d) are weighted by the respective dwell times of each state. This contrasts measurements on graphene, where the fluorescence lifetime of each state is independent of any weighing. As mentioned before, the presented smFRET data are based on the fluorescence intensity and not the fluorescence lifetime as used for GETvNA.

**Fig. S18.**

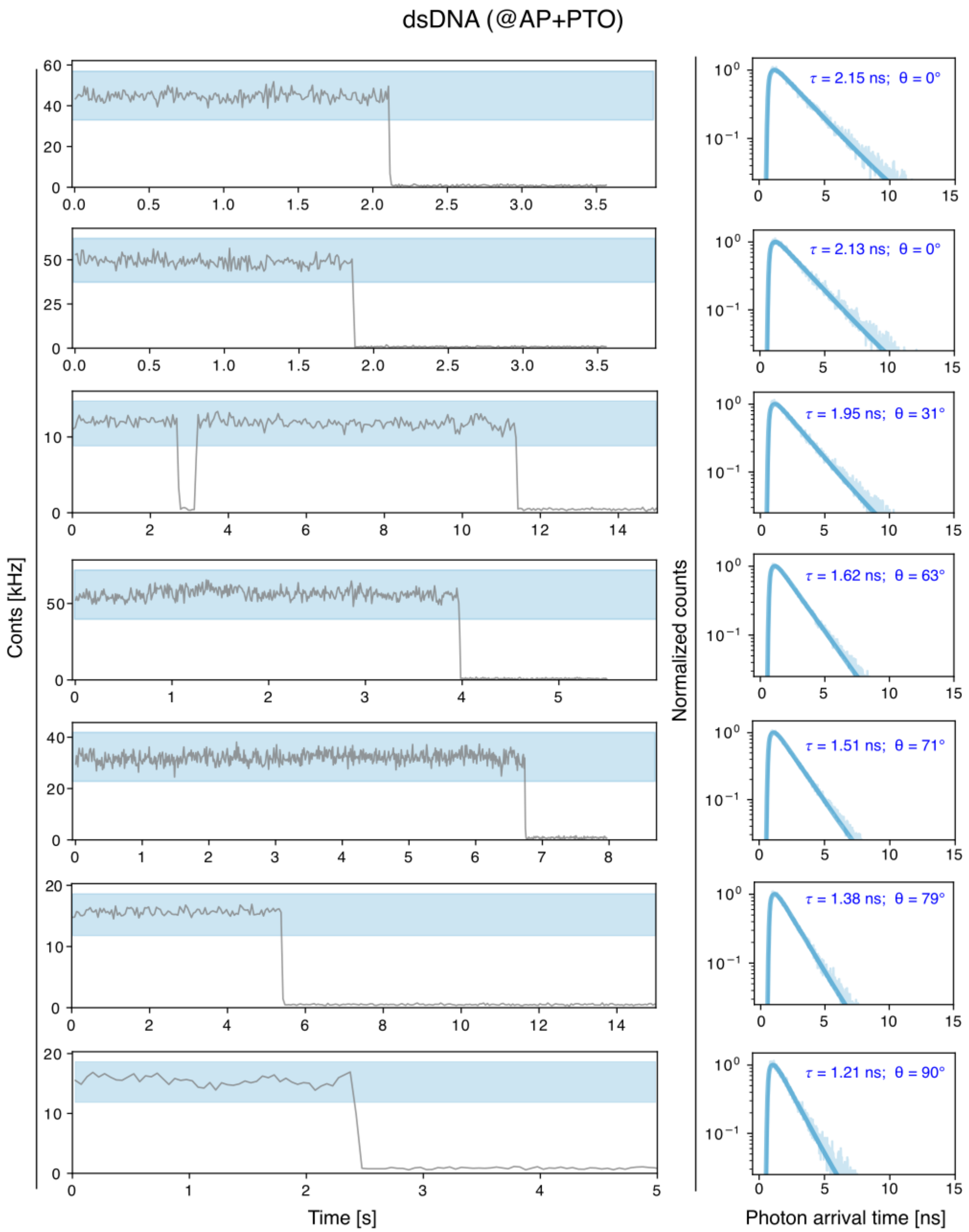

**Fig. S18. Exemplary time traces of systems where the dsDNA segment contained an AP site and a PTO modification, in the absence of Endo IV.** Seven example time traces are shown. The fluorescence intensity time traces are shown on the left, and the fluorescence decays and corresponding monoexponential fits on the right. For each case, the fitted fluorescence lifetime and the corresponding bending angle are shown next to the fluorescence decays. The rectangles from the intensity time traces highlight the photons used to obtain the fluorescence lifetime decay curve.

**Fig. S19.**

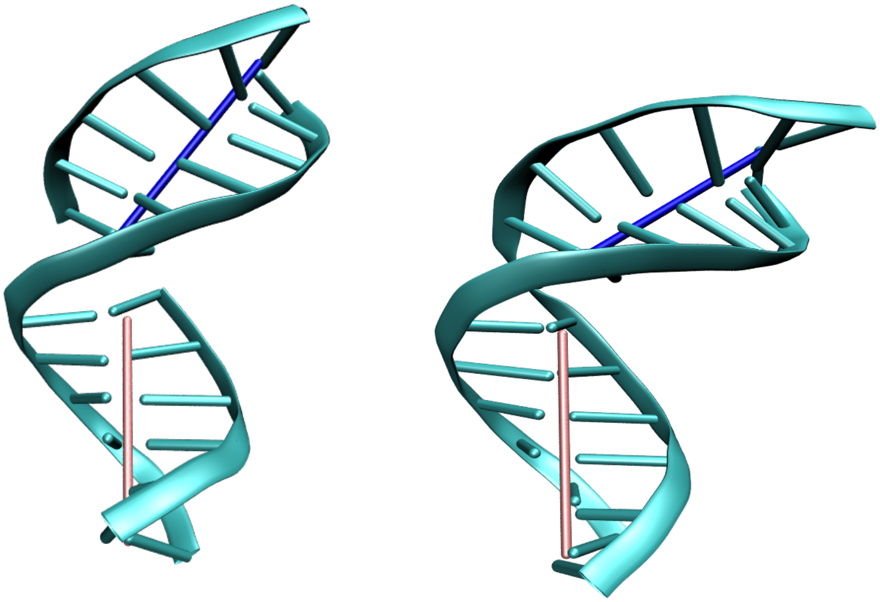

**Fig. S19. Averaged structure of the alpha (left) and beta (right) anomers of dsDNA bearing an AP site.** The shown structures were reconstructed from NMR studies. Cartoon representations of the DNA from the PDB structures 2HSL (alpha) and 2HSS (beta) are shown in pale blue. Superimposed are the two axes of the helix of the two segments divided by the AP site. The angles between both helical axes are 42.3° (alpha) and 53.3° (beta).

**Fig. S20.**

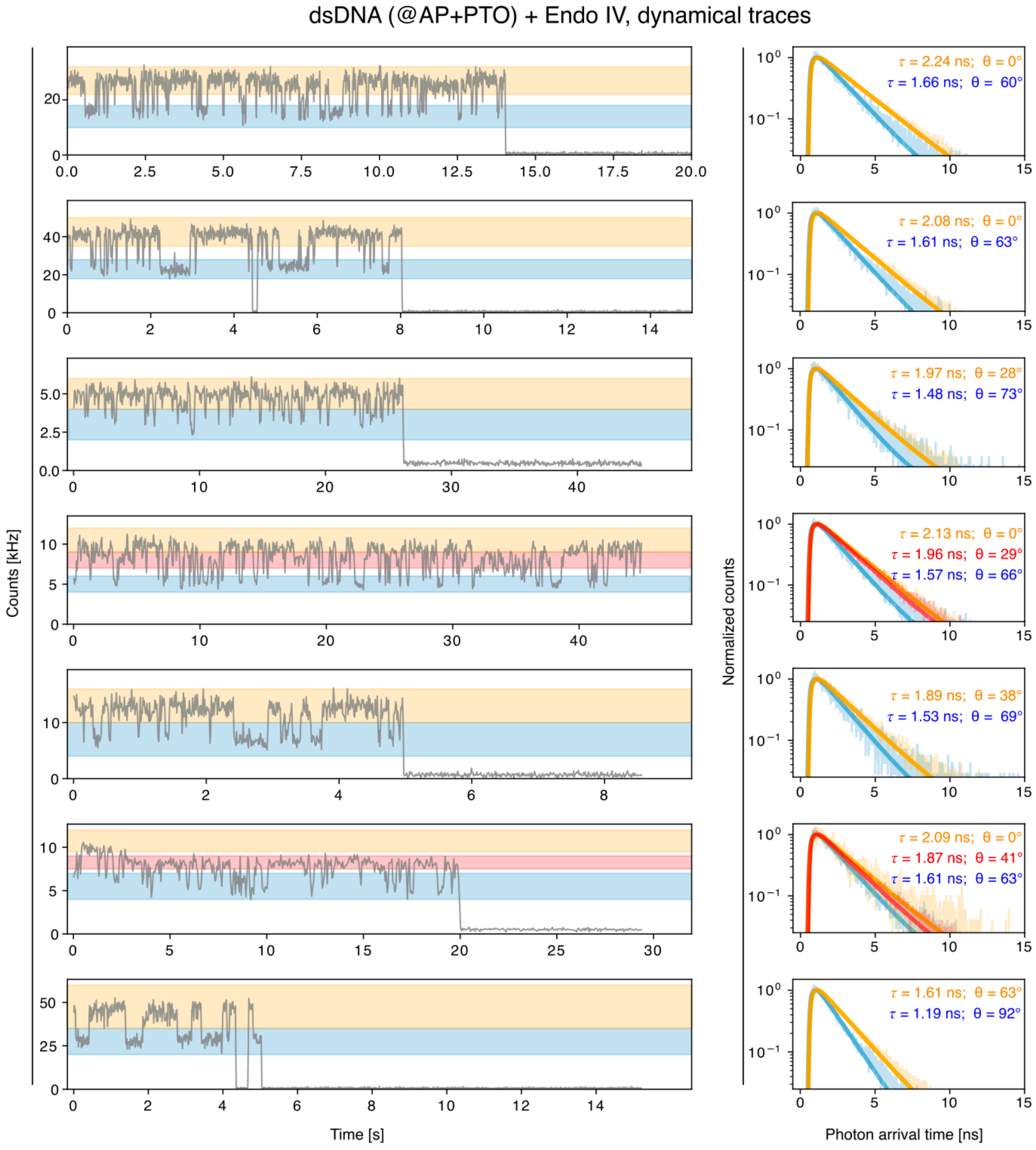

**Fig. S20. Exemplary time traces showing switching between states, corresponding to systems where the dsDNA segment contained an AP site and a PTO modification, in the presence of Endo IV.** Seven exemplary time traces are shown. The fluorescence intensity time traces are depicted on the left, and the fluorescence decays and corresponding monoexponential fits on the right. The color-coded rectangles from the intensity time traces highlight the photons used to obtain each fluorescence lifetime decay curve. The fitted fluorescence lifetime and the corresponding bending angle are shown next to the fluorescence decays.

**Fig. S21.**

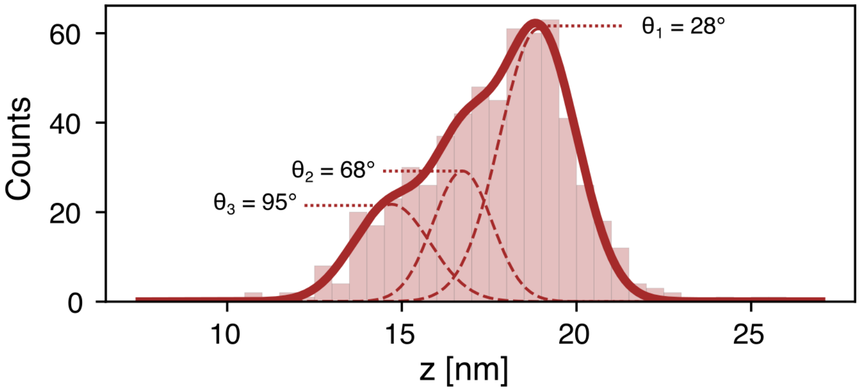

**Fig. S21. Histogram of heights obtained from traces not showing switching between states determined by GETvNA in the presence of Endo IV.** The dsDNA contained an AP site and a PTO modification. For every individual trace, a single height was computed, obtained from the fluorescence lifetime fitted using all the photons detected before photobleaching. A three-peak Gaussian distribution was used to fit the experimental distribution. The bending angles corresponding to each subpopulation are also shown.

**Fig. S22.**

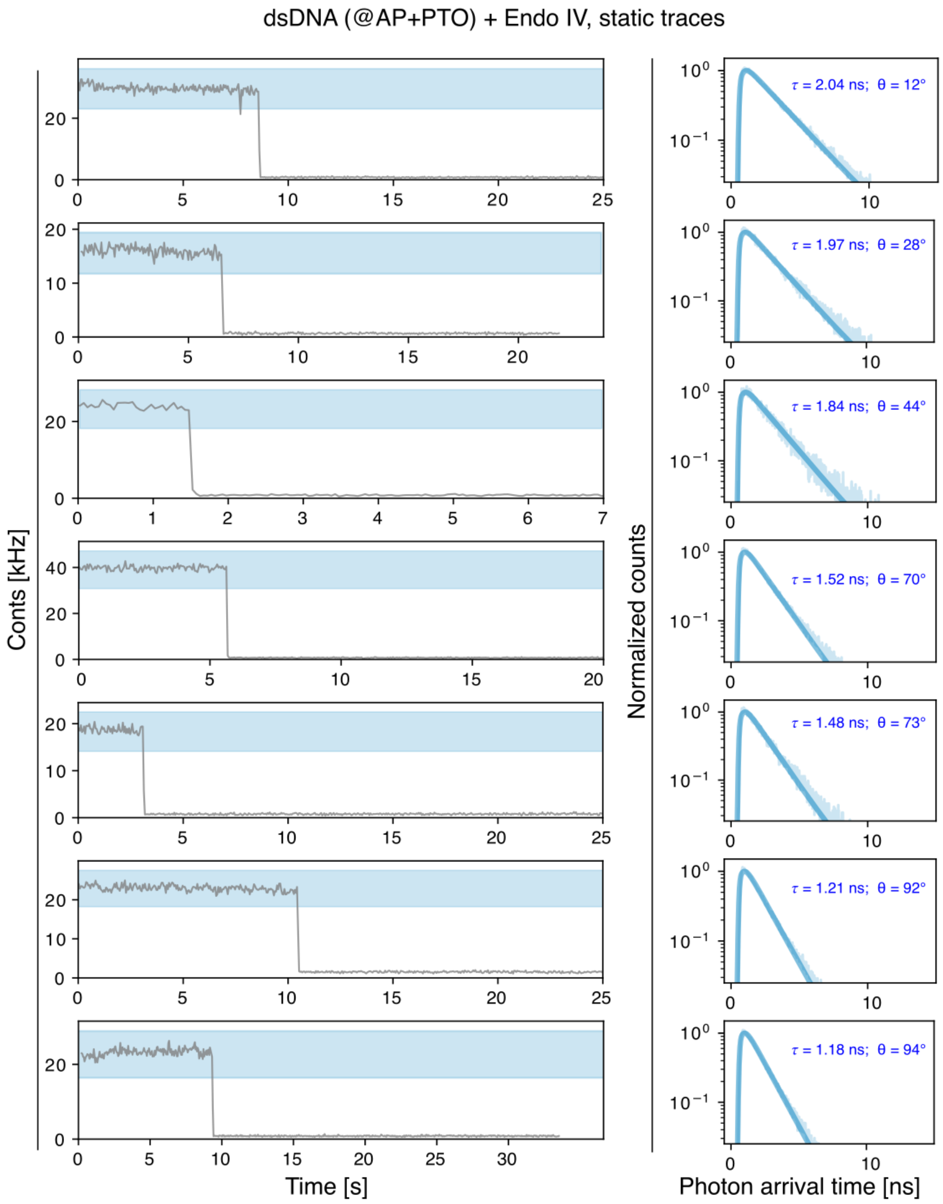

**Fig. S22. Exemplary time traces not exhibiting a state switching in the presence of Endo IV.** Here, the measurements correspond to systems where the dsDNA segment contained an AP site and a PTO modification. Seven example time traces are shown. The fluorescence intensity time traces are depicted on the left, and the fluorescence decays and corresponding monoexponential fits on the right. The rectangles from the intensity time traces highlight the photons used to obtain the fluorescence lifetime decay curves. The fitted fluorescence lifetime and the corresponding bending angle are shown next to the fluorescence decays.

**Fig. S23.**

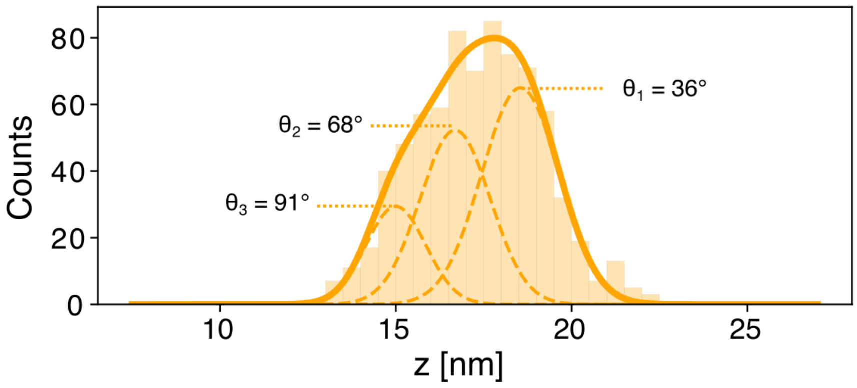

**Fig. S23. Histogram of heights obtained by GETvNA for dsDNA containing an AP site and lacking a PTO modification, in the presence of Endo IV.** For every individual trace, a single height was computed, obtained from the fluorescence lifetime fitted using all the photons detected before photobleaching. A three-peak Gaussian distribution was used to fit the experimental distribution. The bending angles corresponding to each subpopulation are also shown.

**Fig. S24.**

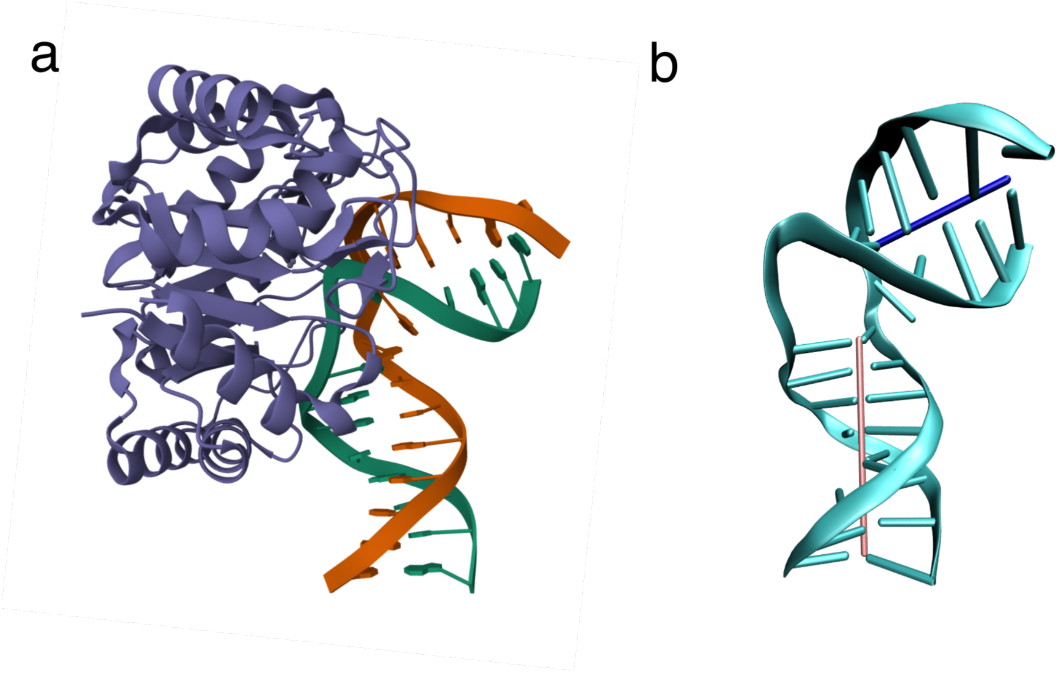

**Fig. S24. Crystal structure of Endo IV bending dsDNA. a)** PDB structure 2NQJ[3] showing Endo IV (purple) recognizing an AP site and introducing subsequent bending of the dsDNA segment. **b)** Helical axis of the Endo IV-bent DNA. A cartoon representation of the DNA in the PDB structure 2NQJ is shown in pale blue. Here, the protein is not shown for clarity. Superimposed are the two axes of the helix (dark blue and red) of the two segments divided by the AP site. The angle between both helical axes is 74.4°.

**Fig. S25.**

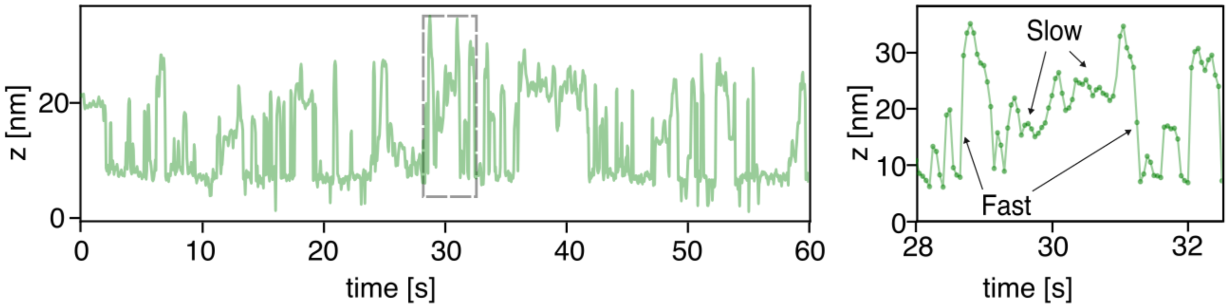

**Fig. S25. Exemplary time trace of AGT cluster diffusing on DNA without A-tract.** Left: 60 second time window. Right: Zoom-in on the region marked by the gray dotted-rectangle. Fast and slow modes are highlighted by arrows.

**Fig. S26.**

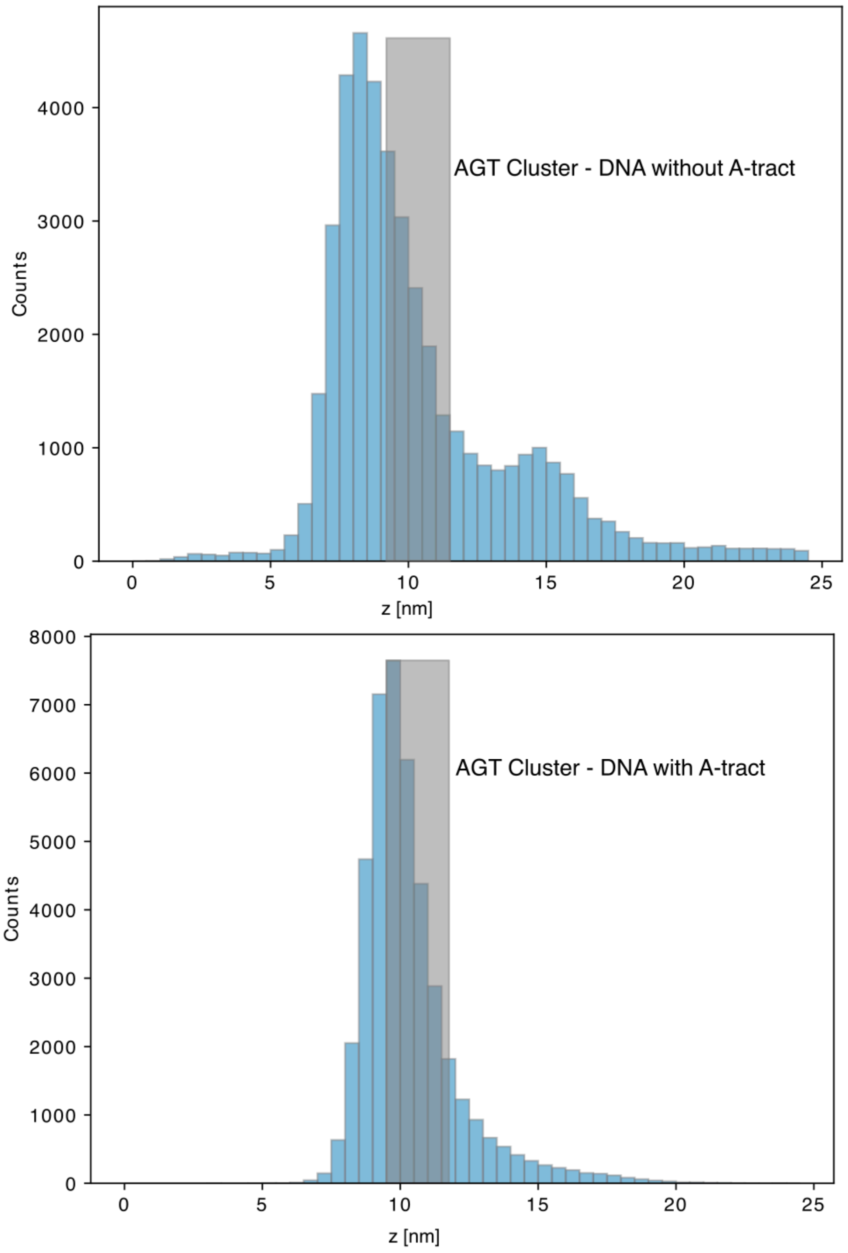

**Fig. S26. Height histograms for AGT cluster, in the absence (top) and in the presence (bottom) of an A-tract.** The gray shaded area represents the height where the A-tract is located (in those samples containing A-tract). For each distribution, twelve traces lasting 180 seconds were used. Here, it should be noticed that cluster growth was produced by the binding of unlabeled AGT to fluorescently labeled AGT already bound to DNA. This explains why the labeled proteins remained around the A-tract region and were not axially shifted in the bottom histogram.

**Fig. S27.**

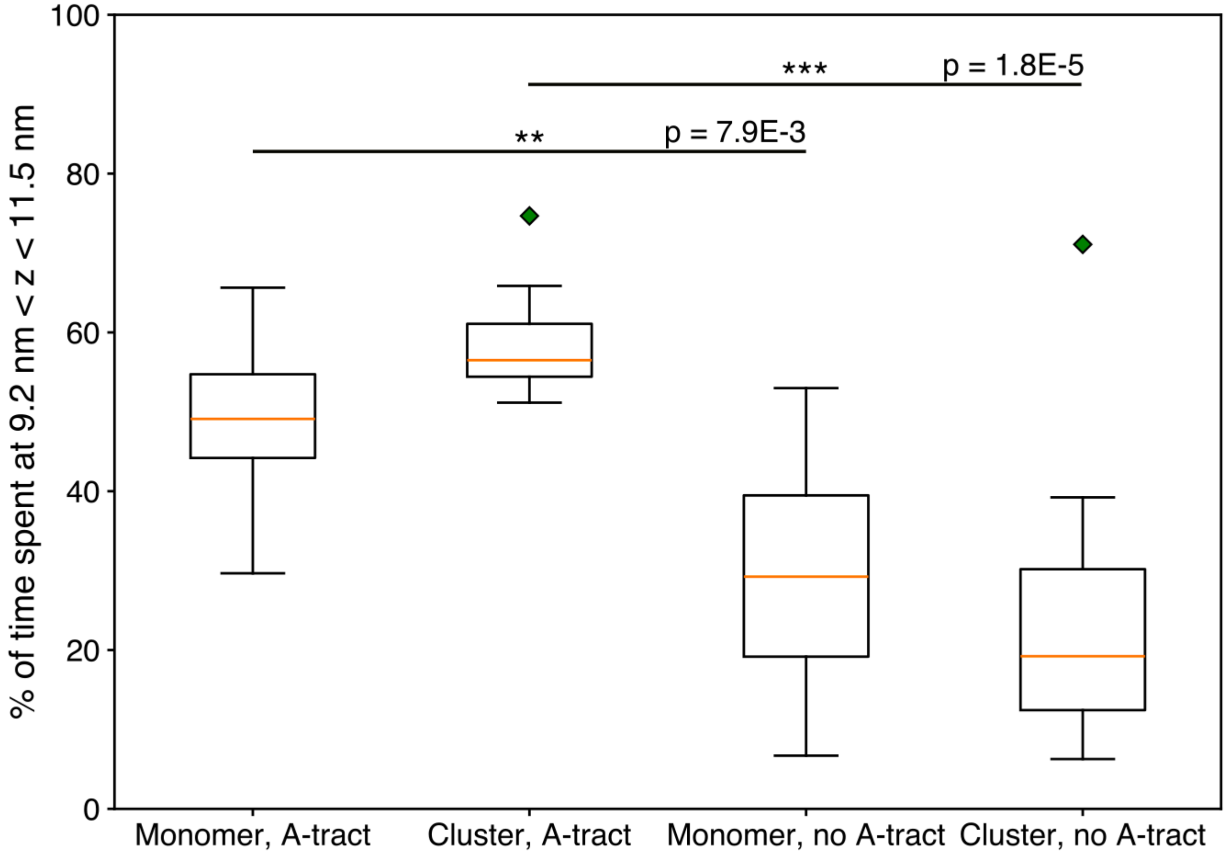

**Fig. S27. Box plot showing the fraction of time spent within the region where the A-tract is located, in samples containing an A-tract.** In particular, AGT clusters showed significantly different residence times at the A-tract region for DNA containing an A- tract compared to DNA without an A-tract. For each condition, twelve traces lasting 180 seconds were used, and from each of them the time spent between 9.2 nm and 11.5 nm was computed. p-values were obtained using a Kolmogorov-Smirnov test to compare the different distributions.

**Fig. S28.**

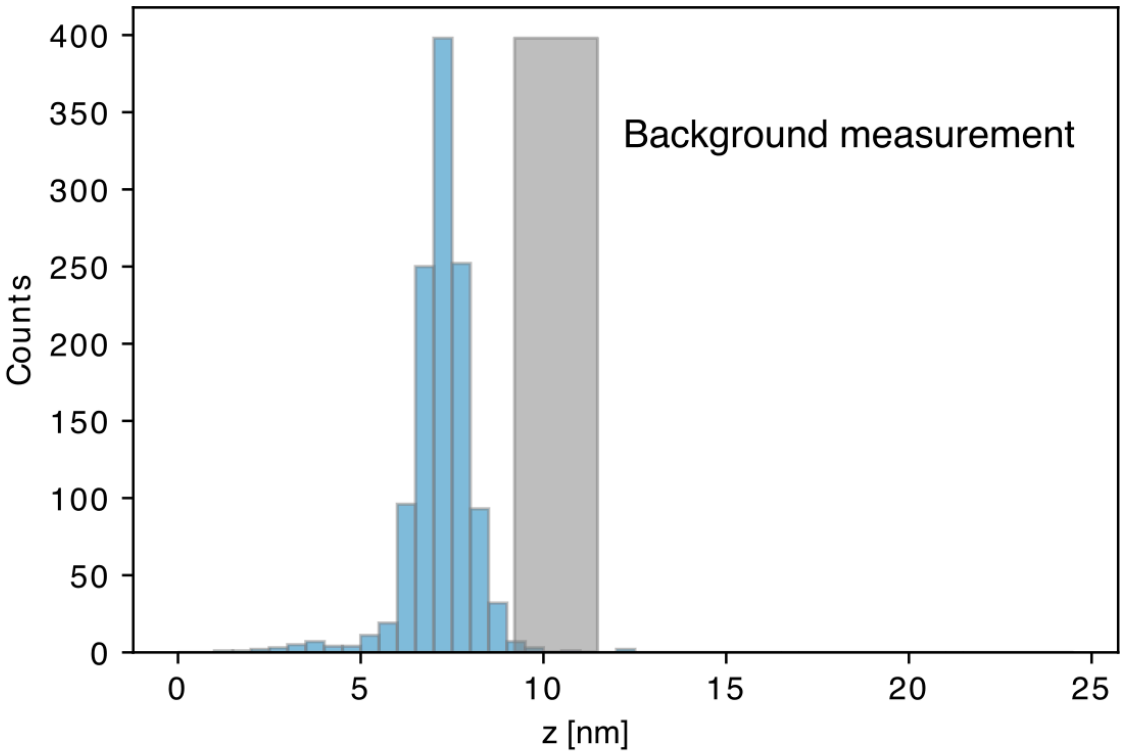

**Fig. S28. Height histogram obtained from traces acquired in background regions in a sample with the DNA construct without an A-tract.** Here, locations were selected, where no specific signal from ATTO 647N was detected. As a visual reference, the gray shaded area represents the height where the A-tract is located in those samples containing A-tract. A mean height of $z$ = 7.5 nm with $\sigma_z$ = 0.9 nm is observed, originating from noisy measurements of fluorescence lifetimes < 0.1 ns.

**Fig. S29.**

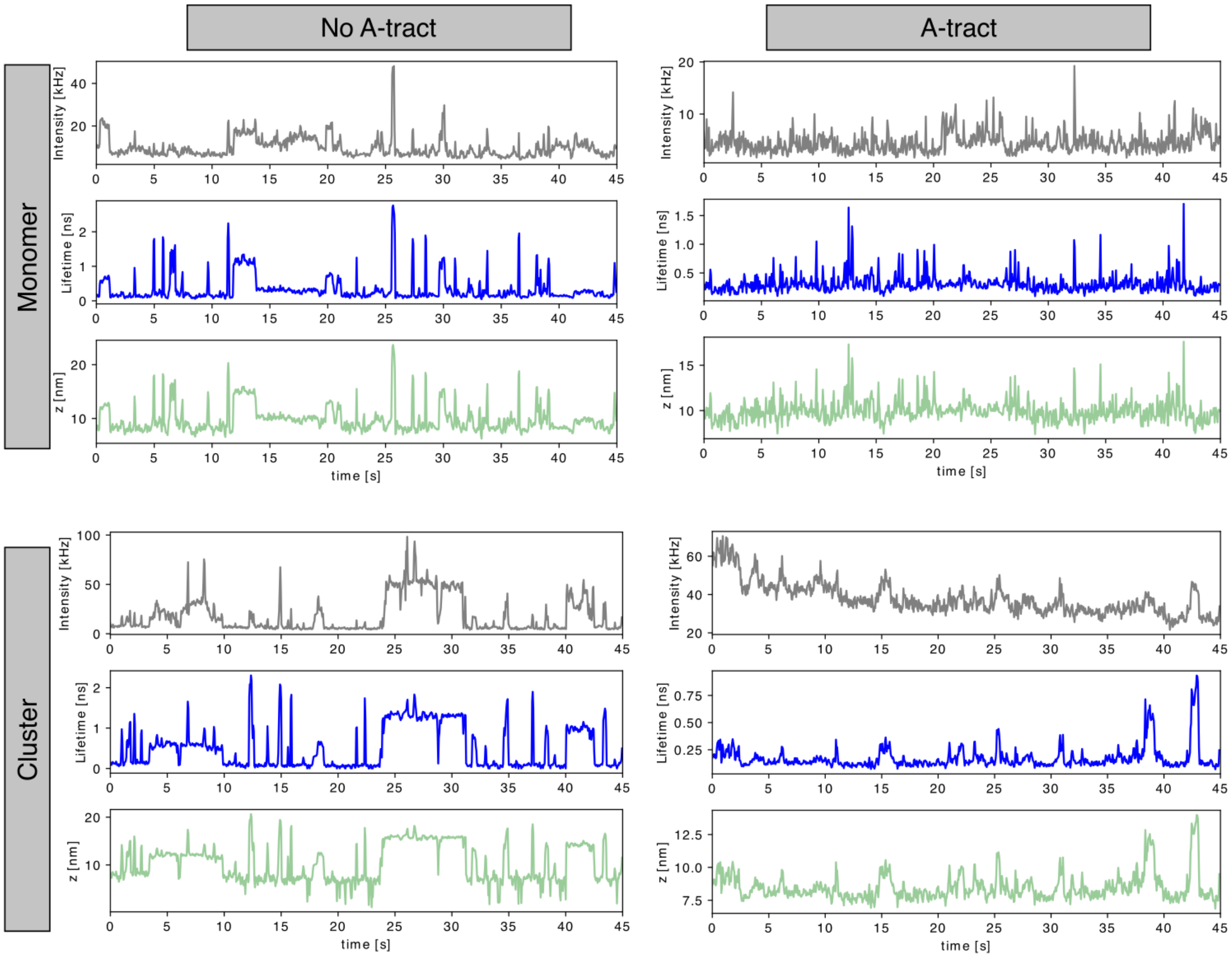

**Fig. S29. Exemplary time traces of AGT clusters and monomers, diffusing along dsDNA containing or lacking A-tract.** For all cases, the fluorescence intensity, fluorescence lifetime and height time traces are depicted.

**Fig. S30.**

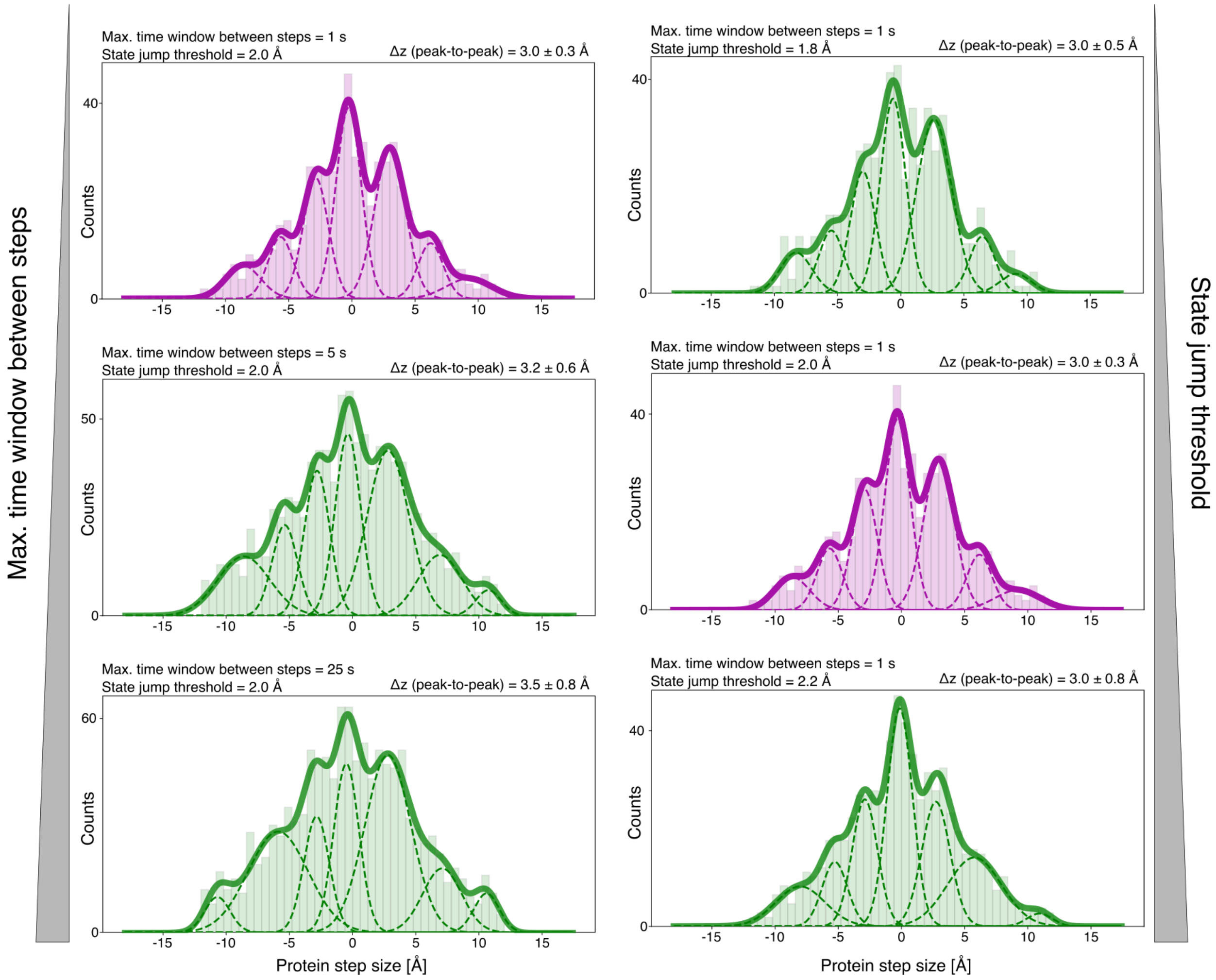

**Fig. S30. Influence of the parameters used to filter and study AGT steps when diffusing on dsDNA containing an A-tract.** The histograms of the protein step size obtained for all the data with DNA after performing the filtering step with different parameters are shown. On the left, the influence of the time window tolerance is depicted, while on the right we present how the height threshold used to detect a protein step affects the step size distribution. For each distribution, a seven-peak Gaussian fit is shown with a solid line, while each component of the fit is shown with a dashed line. The weighted averages of peak-to-peak distances are also presented, together with the weighted standard deviation (weights are calculated from the areas under each Gaussian curve). Whereas the green plots show the tested combination of parameters, the two purple plots represent the final values used in Fig. 4 (main).

**Fig. S31.**

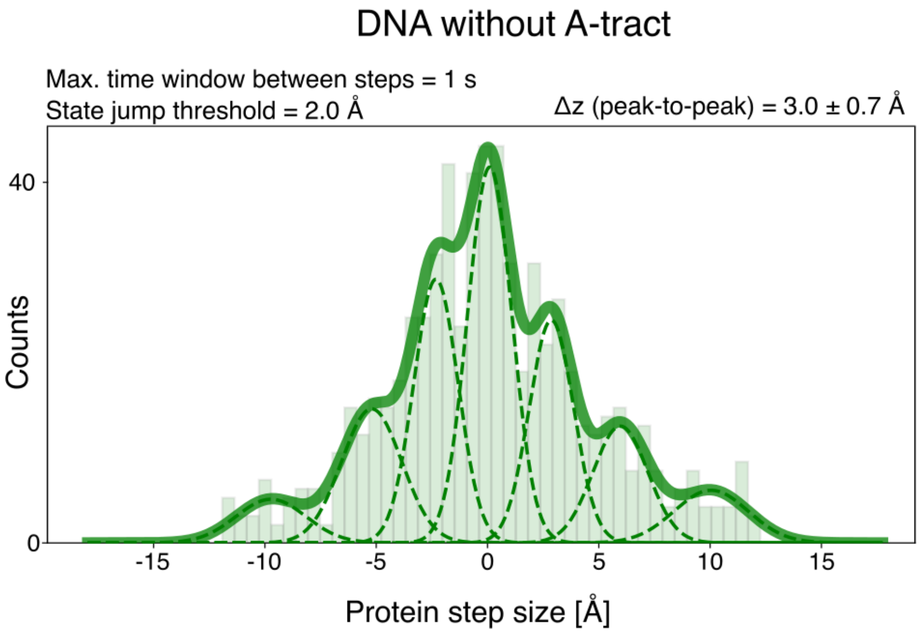

**Fig. S31. Histogram of protein step size, obtained for all the data acquired on DNA lacking an A-tract after performing the filtering step described in section 2.3**. A seven-peak Gaussian fit is shown as a solid line, while each component of the fit is shown with a dashed line. The weighted average of peak-to-peak distances is also presented, together with the weighted standard deviation (weights are calculated from the areas under each Gaussian curve).

**Fig. S32.**

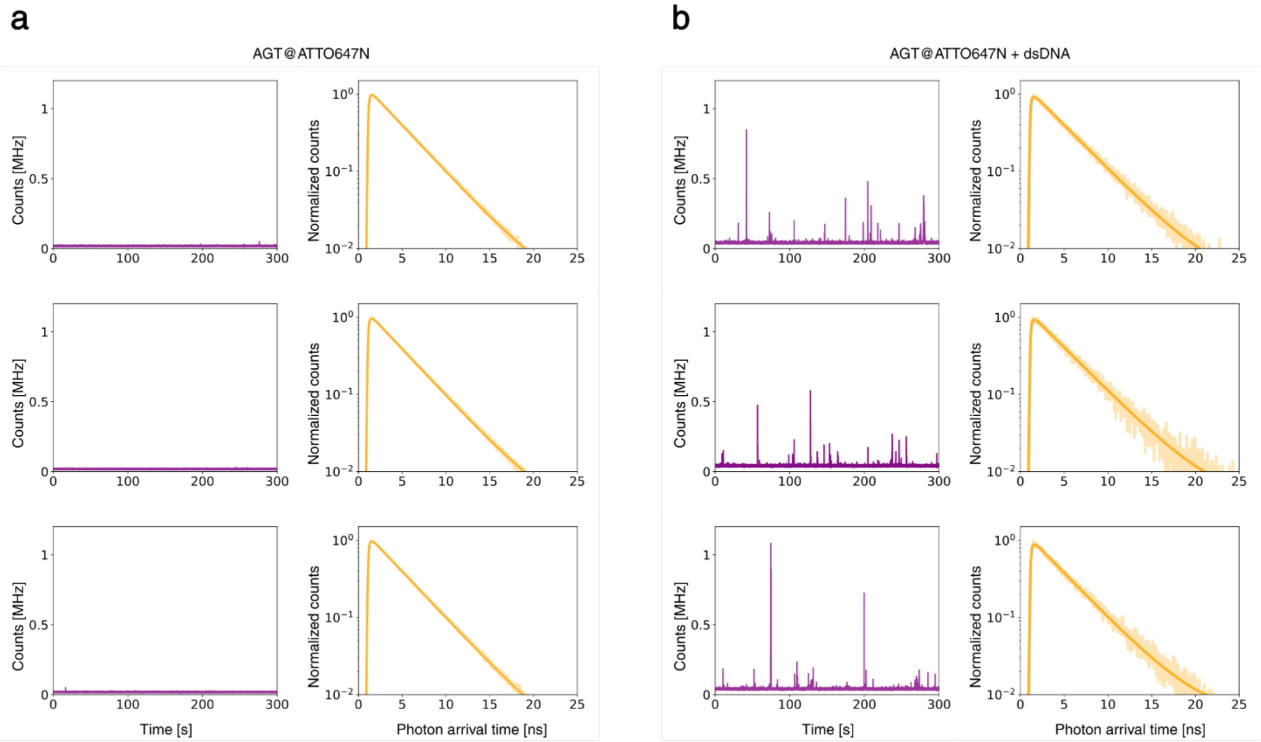

**Fig. S32. Measurement of the reference unquenched lifetime for ATTO647N labeled AGT enzyme bound to dsDNA.** a) Time traces and decay curves relative to 300nM ATTO647N labeled AGT and 3.7 µM unlabeled AGT diluted in AGT experiment buffer and measured in solution with a 640 nm excitation laser. The decay curves are obtained by considering all photons and correspond to an average fluorescent lifetime of 3.58 ns. b) Time traces and decay curves relative to 300 nM ATTO647N labeled AGT, 3.7 µM unlabeled AGT and 10 µM dsDNA (unlabeled 66mer A-tract + 136mer A-tract) diluted in AGT experiment buffer and measured in solution. Intensity spikes appear with the addition of dsDNA and are hence interpreted as DNA-protein complexes diffusing in solution. The decay curves are obtained considering only the photons in the intensity spikes (with a 120 kHz lower threshold in order to have a minimum SBR of 2) and correspond to an average fluorescent lifetime of 3.78 ns, which is then used as the reference unquenched lifetime for AGT measurements.

### 4 Supplementary Tables

#### Table S1.

| Short description | Sequence (5’ to 3’ end) | Supplier | Use |
|---|---|---|---|
| 36mer ATTO 542 | [ATTO542]TAACCACCGTGTACTCGT TATTCGATCCGATCAGTC | Biomers | Labeled strand for 36 bp standing dsDNA (Fig. 1) |
| 37mer ATTO 542 | [ATTO542]TAACCACCGTGTACTCGT TATTCGATCCGATCAGTCC | Biomers | Labeled strand for 37 bp standing dsDNA (Fig. 1) |
| 45mer ATTO 542 | [ATTO542]TAACCACCGTGTACTCGT TATTCGATCCGATCAGTCCGTCGTA GC | Biomers | Labeled strand for 45 bp standing dsDNA (Fig. 1) |
| 51mer ATTO 542 | [ATTO542]TAACCACCGTGTACTCGT TATTCGATCCGATCAGTCCGTCGTA GCCAGTTC | Biomers | Labeled strand for 51 bp standing dsDNA (Fig. 1) |
| 66mer ATTO 542 | [ATTO542]TAACCACCGTGTACTCGT TATTCGATCCGATCAGTCCGTCGTA GCCAGTTCTTAAGACTGTAGATT | Biomers | Labeled strand for 66 bp standing dsDNA (Fig. 1); labeled strand for 3A, 5A and 7A bulged dsDNA (Fig. 2) |
| 136mer | ACGGTCGTCAAATCTGTCGTGGTAG GATCTGTCGTGGTAGGATCTGTCGT GGTAACTTTGGGGTTGGATAAATCT ACAGTCTTAAGAACTGGCTACGACG GACTGATCGGATCGAATAACGAGTA CACGGTGGTTA | IDT | Complementary unlabeled strand for 45 bp, 51 bp and 66 bp unmodified standing dsDNA (Fig. 1); complementary unlabeled strand for abasic site dsDNA, for Endo IV measurements (Fig. 3); complementary unlabeled strand for 66 bp dsDNA without A-tract, for AGT measurements (Fig. 4) |

| Short description | Sequence (5' to 3' end) | Supplier | Use |
|---|---|---|---|
| 73mer | GGGATCACGAAGCTCGAAGCACGC AGTTACCACGACGGACTGATCGGAT CGAATAACGAGTACACGGTGGTTA | IDT | Complementary unlabeled strand for 36 bp and 37 bp unmodified standing dsDNA (Fig. 1) |
| 136mer 3A | ACGGTCGTCAAATCTGTCGTGGTAG GATCTGTCGTGGTAGGATCTGTCGT GGTAACTTTGGGGTTGGATAAATCT ACAGTCTTAAGAACTGGCTACGACG GACTGATAAACGGATCGAATAACGA GTACACGGTGGTTA | IDT | Complementary unlabeled strand presenting 3 additional adenines, for 3A bulged dsDNA (Fig. 2) |
| 136mer 5A | ACGGTCGTCAAATCTGTCGTGGTAG GATCTGTCGTGGTAGGATCTGTCGT GGTAACTTTGGGGTTGGATAAATCT ACAGTCTTAAGAACTGGCTACGACG GACTGATAAAAACGGATCGAATAAC GAGTACACGGTGGTTA | IDT | Complementary unlabeled strand presenting 5 additional adenines, for 5A bulged dsDNA (Fig. 2) |
| 136mer 7A | ACGGTCGTCAAATCTGTCGTGGTAG GATCTGTCGTGGTAGGATCTGTCGT GGTAACTTTGGGGTTGGATAAATCT ACAGTCTTAAGAACTGGCTACGACG GACTGATAAAAAAACGGATCGAATA ACGAGTACACGGTGGTTA | IDT | Complementary unlabeled strand presenting 7 additional adenines, for 7A bulged dsDNA (Fig. 2) |
| 51mer A-tract ATTO 542 | [ATTO542]TAACCACCGTGTACTCGT TATTCGATCCGATCAGTCAAAAAAAG CCAGTTCTTAAGACTGTAGATT | IDT | Labeled strand for 51 bp standing dsDNA with an A-tract (Fig. 2) |
| 136mer A-tract | ACGGTCGTCAAATCTGTCGTGGTAG GATCTGTCGTGGTAGGATCTGTCGT GGTAACTTTGGGGTTGGATAAATCT ACAGTCTTAAGAACTGGCTTTTTTTG ACTGATCGGATCGAATAACGAGTAC ACGGTGGTTA | Biomers | Complementary unlabeled strand for 51 bp dsDNA with A-tract (Fig. 2); complementary unlabeled strand for 66 bp dsDNA with A-tract, for AGT measurements (Fig. 4) |

| Short description | Sequence (5' to 3' end) | Supplier | Use |
|---|---|---|---|
| 66mer AP ATTO 542 | [ATTO542]TAACCACCGTGTACTCGTTAT[Spd]CGATCCGATCAGTCCGTCGTAGCCAGTTCTTAAGACTGTAGATT | Biomers | Labeled strand presenting an abasic site for the dsDNA used for Endo IV measurements (SI) |
| 66mer AP PTO ATTO 542 | [ATTO542]TAACCACCGTGTACTCGTTAT*[Spd]CGATCCGATCAGTCCGTCGTAGCCAGTTCTTAAGACTGTAGATT | Biomers | Labeled strand presenting an abasic site and an adjacent PTO modification for the dsDNA used for Endo IV measurements (Fig. 3) |
| 66mer internal ATTO 542 | TAACCACCGTGTACTCGTTAT[dT-ATTO542]CGATCCGATCAGTCCGTCGTAGCCAGTTCTTAAGACTGTAGATT | Biomers | Labeled strand internally labeled at base 45 with ATTO 542, for external-internal labeling comparison (SI) |
| 66mer internal ATTO647N | TAACCACCGTGTACTCGTTATTCGATCCGATCAGTCCG[dT-ATTO647N]CGTAGCCAGTTCTTAAGACTGTAGATT | Biomers | Labeled strand internally labeled at base 28 with ATTO647N, for A-tract height measurement (SI) |
| Red-labeled strand for smFRET | [Alexa647]CGAGAAGATTCAAGGTTCAC | Eurofins | Alexa 647-labeled strand for the dsDNA used in smFRET Endo IV measurements (SI) |
| Green-labeled strand for smFRET | AACGACGGACAGTGACAAGC[Cy3] | Eurofins | Cy3-labeled strand for the dsDNA used in smFRET Endo IV measurements (SI) |
| AP PTO strand for smFRET | GTGTGATG*[Spd]AGGTGGTG | Eurofins | Strand presenting an abasic site and an adjacent PTO modification for the dsDNA used in smFRET Endo IV measurements (SI) |
| AP strand for smFRET | GTGTGATG[Spd]AGGTGGTG | IDT | Strand presenting an abasic site for the dsDNA used in |

| Short description | Sequence (5' to 3' end) | Supplier | Use |
|---|---|---|---|
| | | | smFRET Endo IV measurements (SI) |
| Reverse complementary with bioteg for smFRET | GTGAACCTTGAATCTTCTCGCACCACCTACATCACACGCTTGTCACTGTCCGTCGTT[Bioteg] | Eurofins | Complementary unlabeled strand used in smFRET study (SI) |
| Biotinylated 36mer | [Biotin]TTGACTGATCGGATCGAATAACGAGTACACGGTGGTTA | Biomers | Complementary unlabeled strand used for solution measurements (SI) |
| 66mer A-tract Cy3B | [Cy3B]TAACCACCGTGTACTCGTTATTCGATCCGATCAGTCAAAAAAAGCCAGTTCTTAAGACTGTAGATT | Eurofins | Labeled strand for 66 bp standing dsDNA with an A-tract, for AGT measurements (Fig. 4) |
| 66mer Cy3B | [Cy3B]TAACCACCGTGTACTCGTTATTCGATCCGATCAGTCCGTCGTAGCCAGTTCTTAAGACTGTAGATT | Eurofins | Labeled strand for 66 bp standing dsDNA without an A-tract, for AGT measurements (Fig. 4) |
| Unlabeled 66mer A-tract | TAACCACCGTGTACTCGTTATTCGATCCGATCAGTCAAAAAAAGCCAGTTCTTAAGACTGTAGATT | Eurofins | Unlabeled strand for dsDNA in solution for measurement of unquenched lifetime of ATTO647N for AGT measurements (SI) |

List of DNA oligonucleotides used within this study.

**Table S2.**

| Quantity | Value |
|---|---|
| $d_{0,\ ATTO\ 542}$ | 17.7 nm |
| $d_{0,\ ATTO\ 647N}$ | 18.5 nm |
| $\tau_{0,\ ATTO\ 542}$ in Tx-Tq-PCA with 1× PCD | 3.51 ns |
| $\tau_{0,\ ATTO\ 542}$ in NEB3-Tx-Tq-PCA with 1× PCD | 3.52 ns |
| $\tau_{0,\ ATTO\ 542}$ internally-labeled in Tx-Tq-PCA with 1× PCD | 3.40 ns |
| $\tau_{0,\ ATTO\ 647N}$ with AGT bound to DNA in AGT experiment buffer | 3.78 ns |
| $\tau_{0,\ ATTO\ 647N}$ internally-labeled in Tx-Tq-PCA with 1× PCD | 3.74 ns |

List of used reference parameters for computing the distance from graphene for a measured fluorescence lifetime and a given fluorophore. The listed fluorescence lifetimes are deconvoluted.
